# Defining and predicting transdiagnostic categories of neurodegenerative disease

**DOI:** 10.1101/664250

**Authors:** Eli J. Cornblath, John L. Robinson, Virginia M.-Y. Lee, John Q. Trojanowski, Danielle S. Bassett

## Abstract

In current models of neurodegeneration, individual diseases are defined by the presence of one or two pathogenic protein species. Yet, it is the rule rather than the exception that a patient meets criteria for more than one disease. This fact often remains hidden until autopsy, when neuropathological evaluation can assign disease labels based on gold-standard criteria. Ultimately, the prevalence of concomitant diagnoses and the inability to infer an underlying neuropathological syndrome from clinical variables hinders the identification of patients who might be good candidates for a particular intervention. Here, by applying graph-based clustering to post-mortem histopathological data from 1389 patients with degeneration in the central nervous system, we generate 4 non-overlapping, data-driven disease categories that simultaneously account for amyloid-*β* plaques, tau neurofibrillary tangles, *α*-synuclein inclusions, neuritic plaques, TDP-43 inclusions, angiopathy, neuron loss, and gliosis. The resulting disease clusters are transdiagnostic in the sense that each cluster contains patients belonging to multiple different existing disease diagnoses, who colocalize in clusters according to the pathogenic protein aggregates known to drive each disease. We show that our disease clusters, defined solely by histopathology, separate patients in terms of cognitive phenotypes, cerebrospinal fluid (CSF) protein levels, and genotype in a manner that is not trivially explained by the representation of individual diseases within each cluster. Finally, we use cross-validated multiple logistic regression to generate high accuracy predictions (AUC *>* 0.9) of membership to both existing disease categories and transdiagnostic clusters based on CSF protein levels and genotype, both accessible *in vivo*. Broadly, our approach parses phenotypic and genotypic heterogeneity in neurodegenerative disease, and represents a general framework for identifying otherwise-fuzzy disease subtypes in other areas of medicine, such as epilepsy, vascular disease, and cancer. In clinical neurology, the statistical models we generate may be useful for repurposing drugs by comparing efficacy to probabilistic estimates of disease cluster membership, as well as for future trials that could be targeted towards an algorithmically defined family of diseases.

## INTRODUCTION

Age-related neurodegenerative diseases affect over 7 million Americans^1^, amounting to nearly $1 trillion in healthcare costs annually. This public health issue is projected to worsen^1–3^ as life expectancy increases and the U.S. population continues to skew towards older individuals. The neurodegenerative disease umbrella includes major clinicopathological entities such as Alzheimer’s disease^4^, Parkinson’s disease^5^, and frontotemporal dementia^6^, in addition to less common disease subtypes such as progressive supranuclear palsy^6^, corticobasal degeneration^6^, and multiple systems atrophy^7^. Neurology is in dire need of translational research that accounts for the heterogeneous presentations of neurodegeneration to facilitate the development of targeted treatments.

Decades of evidence support the notion that pathological protein aggregation is a primary disease process in neurodegeneration^8–10^. These protein aggregates may spread along large white matter fibers over time, causing dysfunction in distant regions^11,12^, and their toxicity is thought to be mediated in part by the inflammatory system^13,14^. Different neurodegenerative syndromes are characterized by aggregation of specific proteins; classically, Alzheimer’s disease involves both amyloid-*β* and microtubule-associated protein tau (tau)^8–10^, Parkinson’s disease involves *α*-synuclein^5^, and frontotemporal dementia can involve tau^6,15^ or TDP-43^16^.

Despite this apparent specificity, aggregation of amyloid-*β*, tau, *α*-synuclein, and TDP-43 is found postmortem in virtually all brains with neurodegenerative disease in addition to brains from cognitively healthy individuals^17–20^. To complicate matters further, numerous *in vitro* and animal studies have demonstrated that these proteins interact to produce unique, concomitant dysfunction^21–24^. Moreover, there are multiple mechanisms by which molecular pathology causes cellular dysfunction^25^ (i.e. angiopathy, gliosis, neuronal cell death, signaling dysfunction), and cellular dysfunction may be a more specific marker of cognitive dysfunction than plaque burden alone^26,27^.

The healthcare system is familiar with the problem of highly complex biology underlying variable clinical presentations. The ever-decreasing costs of computing hardware and easily accessible programming libraries for advanced computational approaches, such as machine learning and network science, provide tools to parse heterogeneity by defining new data-driven disease subtypes. These tools have been applied in multiple contexts in cancer biology^28,29^, epilepsy^30,31^, and psychiatry^32–34^. Machine learning techniques and network approaches have been utilized in speech recordings^35^, neuroimaging^36,37^, and clinical data^38^ of patients with neurodegenerative diseases, but have thus far been underutilized in the field of neuropathology, in which multiple forms of pathological protein aggregates with different morphologies can be measured alongside cellular dysfunction. It is also difficult to map specific imaging phenotypes or biomarkers to a particular disease due to inaccuracies in clinical diagnoses^39^. Indeed, the gold standard for identifying a particular neuropathological syndrome is evaluation of proteinopathic burden on autopsy^4,6,40^.

Here, we use basic statistical approaches to analyze copathology between amyloid-*β*-containing plaques, *α*-synuclein inclusions, tau neurofibrillary tangles, TDP-43 inclusions, neuritic plaques, neuronal loss, angiopathy, and gliosis across 18 brain regions in a sample of 1389 patients evaluated by expert neuropathologists on autopsy. Next, we used a graph-based clustering approach that assigns each patient to a single, data-driven, transdiagnostic “disease cluster” while accounting for all available forms of pathology. We evaluated this approach alongside the existing model of neurodegenerative disease, in which diseases are defined by 1-2 protein species and patients simultaneously meet criteria for multiple diagnoses. Consistent with the traditional understanding of neurodegenerative disease entities, the resulting clusters grouped together diseases known to be driven by the same pathogenic protein. We also found that these disease clusters, which are defined solely by histopathology, differed in terms of cognitive phenotype, cerebrospinal fluid (CSF) protein levels, and genotype at the *APOE* and *MAPT* loci. Importantly, differences in CSF protein levels and genotype remained after controlling for the presence of individual diseases, suggesting that our pathology-defined clusters map to transdiagnostic phenotypic and genotypic boundaries. Finally, using multiple logistic regression, we achieved highly accurate identification (AUC *>* 0.9) of both existing disease labels and data-driven disease clusters from a heterogeneous clinical population based solely on data available *in vivo*. Our findings challenge current definitions of neurodegenerative disease syndromes and provide clinicians with greater explanatory power for parsing disease heterogeneity in the context of existing biomarkers.

## RESULTS

### Copathology-driven clusters group heterogeneous diseases by underlying proteinopathies

Neurodegenerative diseases are often characterized by archetypal distributions of a single protein aggregate^42^, yet co-occurring pathology involving additional forms of protein aggregation can be found in many cases^17^. The vast degree of overlap in molecular and cellular pathology across neurodegenerative diseases complicates the process of assigning patients to meaningful disease categories. To address this problem, we sought to identify new categories of neurodegenerative disease that explicitly account for copathology. Here, using a sample of 1389 autopsy cases with various neurodegenerative diagnoses, we computed the Spearman correlation between vectors of pathology scores for each pair of patients as a measure of pairwise similarity across several features of molecular and cellular pathology (Fig. 2a). These “pathological features” included measurements of different types of proteinopathic features (amyloid-*β* antibody-staining plaques, thioflavin-staining neuritic plaques, tau neurofibrillary tangles, *α*-synuclein inclusions, TDP-43 inclusions, ubiquitin) or types of histological features (angiopathy, gliosis, neuron loss) taken from specific brain regions. When patients are ordered by their arbitrary subject number, the resulting matrix of inter-subject Spearman correlations has no obvious structure (Fig. 2a). When patients are ordered by primary histopathological diagnosis, block-like structure becomes evident, indicating similar histopathological findings in patients with the same primary diagnoses (Fig. 2b). However, we also observe large, positive correlation values on the off-diagonal blocks, indicating similarity in histopathological findings that is unexplained by the primary diagnosis.

**FIG. 1.**
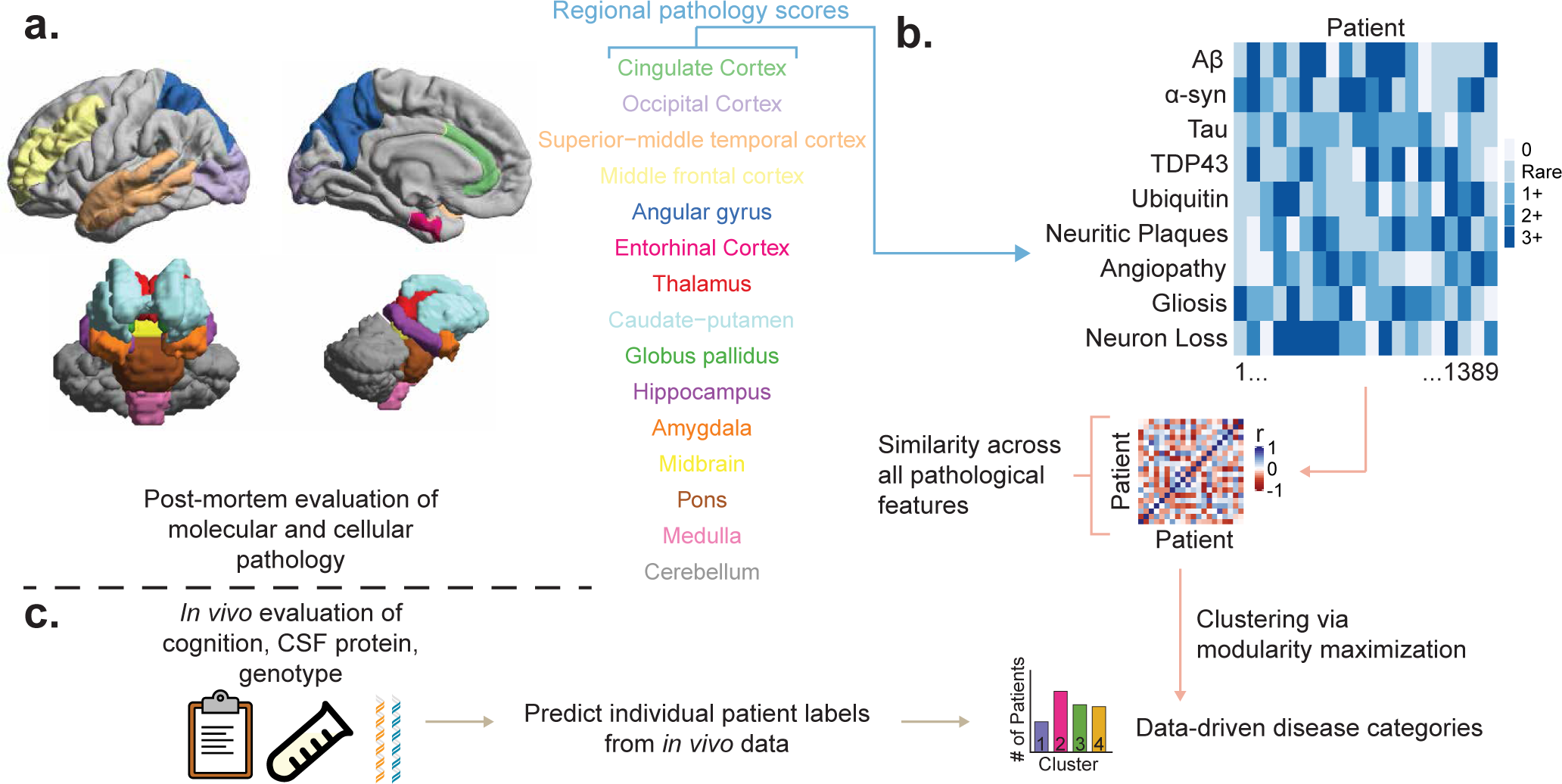
Schematic of data processing. *(a)* The burden of amyloid-*β* plaques, *α*-synuclein plaques, tau neurofibrillary tangles, TDP-43 inclusions, ubiquitin, neuritic plaques, angiopathy, gliosis, and neuron loss was evaluated on a 5-tier ordinal scale (0, Rare, 1+, 2+, 3+) via cerebral autopsy in 1389 patients through the Integrated Neurodegenerative Disease Database^41^. Evaluation of pathological burden was performed for all proteins for the listed regions, in addition to the substantia nigra and locus coeruleus, which are hidden for ease of visualization. Dentate gyrus and CA1/subiculum were quantified separately but shown together here as the hippocampus for ease of visualization. *(b)* We compute a 1389 *×* 1389 similarity matrix whose *ij*^th^ element contains a Spearman correlation between pathology score vectors for patient *i* and patient *j*. Next, we use network community detection by modularity maximization to assign each patient to a data-driven disease cluster. *(c)* Using linked data from cerebrospinal fluid (CSF) protein testing and genotyping, we train statistical models to predict membership to disease clusters.

**FIG. 2.**
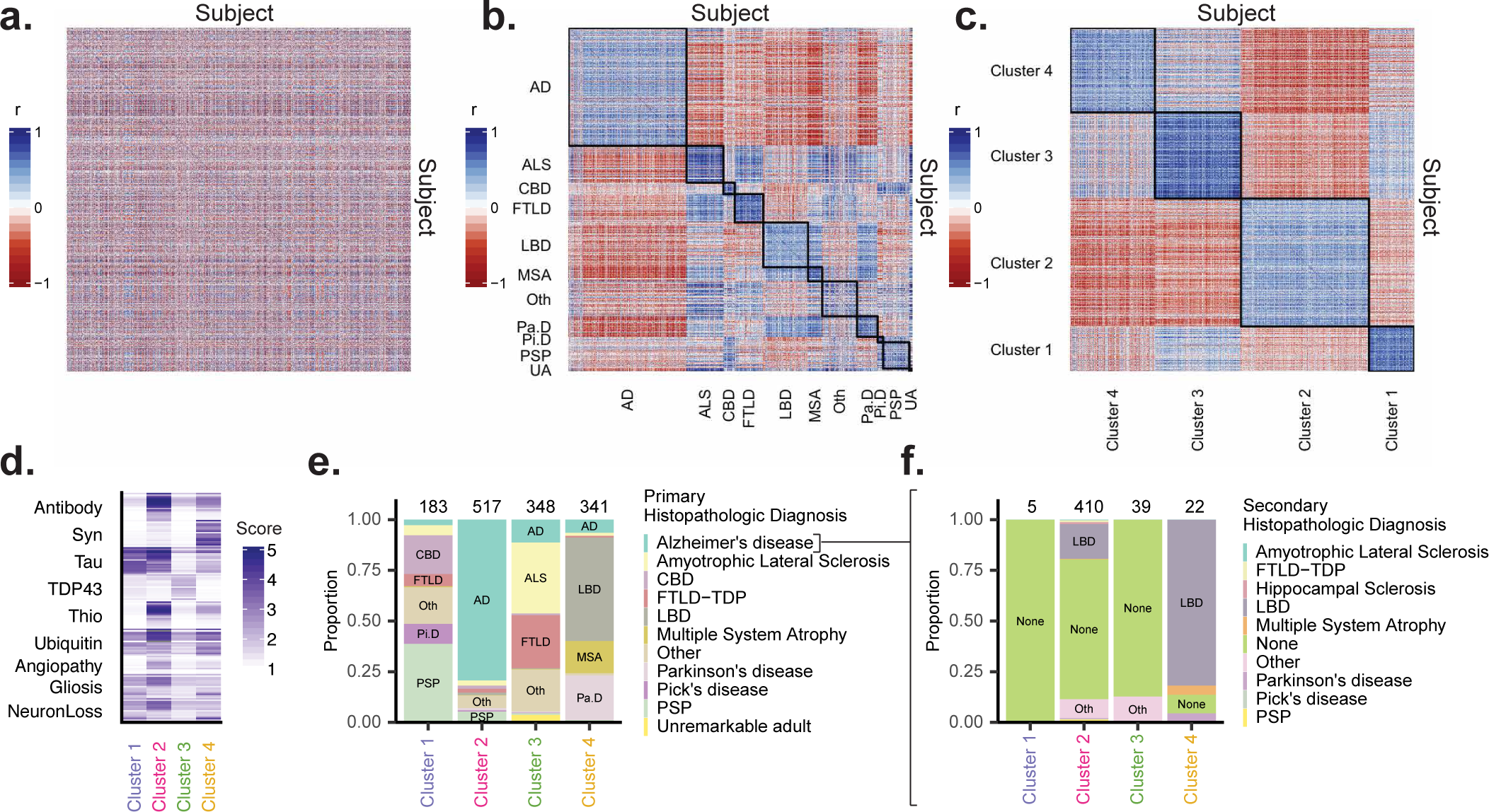
Unsupervised clustering of copathology groups disease entities into proteinopathy families. *(a)* We computed a matrix of Spearman correlations between vectors of pathology scores for each pair of subjects across all available pathological features to quantify the similarity in distributions of pathology. *(b)* The same matrix as in panel *(a)* where rows and columns are ordered by primary histopathologic diagnosis. Black lines along the diagonal mark blocks of patients with the same diagnosis. *(c)* The same matrix as in panel *(a)* where rows and columns are ordered by communities detected through modularity maximization^43^. Black lines along the diagonal mark blocks of patients grouped into the same cluster. *(d)* Representative vector of pathology scores for each cluster (cluster centroids) demonstrate distinct profiles of pathology that map to underlying molecular drivers of disease, including tau, amyloid-*β*, TDP-43, and *α*-synuclein. *(e)* Composition of each cluster in terms of primary histopathologic diagnoses. Each cluster is comprised of disease entities that are putatively caused by the protein most highly represented in the cluster’s centroid. Counts placed above stacked bars indicate the number of patients in each cluster. *(f)* In a subset of patients, all of which have a primary diagnosis of Alzheimer’s disease, we show the composition of each cluster in terms of secondary histopathologic diagnosis. Counts placed above stacked bars indicate the number of patients with Alzheimer’s disease in each cluster. CBD = corticobasal degeneration, FTLD = frontotemporal lobar dementia with TDP-43 inclusions, LBD = dementia with Lewy bodies, PSP = progressive supranuclear palsy.

To parse the observed overlap between disease entities, we grouped patients according to their distributions of pathology using an unsupervised community detection method for networks, known as modularity maximization^43^ (Fig. 2c, see Methods and Supporting Information for details). Notably, this algorithm is agnostic to histopathologic diagnosis, yet consistently grouped histopathologic diagnoses together by their underlying molecular drivers. This fact became evident when we constructed a representative patient for each cluster by calculating the average pathology scores across all patients in that cluster. Specifically, we saw clusters characterized by tau (Cluster 1), amyloid=*β* (Cluster 2), TDP-43 (Cluster 3), and *α*-synuclein (Cluster 4) (Fig. 2d). Indeed, the subjects belonging to Cluster 1 were composed of tauopathy-family frontotemporal lobar dementia (FTLD) syndromes^6^, such as Pick’s Disease, progressive supranuclear palsy, and corticobasal degeneration (Fig. 2e). Cluster 2 was composed primarily of patients with a diagnosis of Alzheimer’s disease (Fig. 2e). Cluster 3 exhibited strong representation of TDP-43 proteinopathies, namely FTLD-TDP and amyotrophic lateral sclerosis (ALS) (Fig. 2e). Finally, Cluster 4 contained primarily synucleinopathies, housing patients with Lewy Body Dementia (LBD), Parkinson’s Disease, and Multiple Systems Atrophy (Fig. 2e). These findings suggest that the solution of our clustering algorithm respects the known hierarchy of neurodegenerative diseases, which is driven by aggregation of specific pathogenic proteins.

After establishing the similarity between our clustering solution and existing schema for categorizing neurodegenerative disease, we next explored how our clustering solution might expand upon these schema. Notably, every cluster contained patients with a primary histopathologic diagnosis of Alzheimer’s disease (Fig. 2e). To understand how our algorithm separated patients within Alzheimer’s disease, we visualized the distribution of secondary histopathologic diagnoses in each cluster in a subset of patients with primary diagnoses of Alzheimer’s disease (Fig. 2f). This analysis revealed that Alzheimer’s patients in the tauopathy and TDP-43 proteinopathy clusters largely did not have any additional diagnoses, while most individuals in the synucleinopathy cluster had LBD as a secondary diagnosis (Fig. 2f). Interestingly, Cluster 2 mostly contained individuals that did not have any secondary diagnoses, but also contained a smaller cohort of individuals that did have a secondary LBD diagnoses, suggesting that the algorithm identified two subgroups of patients with concomitant amyloid-*β*, tau, and *α*-synuclein pathology.

In order to ascertain whether global levels of each protein or specific regional distributions of each protein were primarily driving the algorithm’s separation of patients into clusters, we plotted the mean pathology scores for each cluster (Fig. 2d) on a spatial map of the brain using neuroimaging tools^44,45^ (Fig. S3a). This visualization suggests that differences between clusters are primarily driven by global levels of 1 to 3 pathological features, although Cluster 1 and Cluster 2 differed slightly in their regional distributions of tau neurofibrillary tangles despite both having relatively high tau scores globally. Cluster 1 exhibited greater tau burden uniformly throughout the subcortex while Cluster 2 exhibited low tau in the cerebellum and high tau in the amygdala (Fig. S3a). Overall, these findings show that simultaneously accounting for multiple forms of pathology produces disease labels that both expand and respect the boundaries of existing diagnoses.

### Disease clusters exhibit unique *in vivo* phenotypes

After generating new disease categories based solely on post-mortem pathology, we sought to characterize patients within each cluster in terms of *in vivo* phenotypes. First, we evaluated cognition in an *n* = 147 subsample of patients with available data from the Montreal Cognitive Assessment (MoCA)^46^. To compare MoCA scores between clusters, we defined *M*_*i-j*_ as the median difference between scores from patients in cluster *i* and scores from patients in cluster *j*. Because scores were not all normally distributed, we used the Wilcoxon rank-sum test to evaluate the null hypothesis that *M*_*i-j*_ = 0 for each MoCA subscore and all unique pairs *ij* where *i ≠ j*. We corrected for multiple comparisons across all 6 unique pairwise comparisons for all 6 MoCA subscores by adjusting the false discovery rate (*q <* 0.05). For repetition, naming, orientation, and delayed recall subsections, Cluster 2 had lower median scores than at least one other cluster, with the largest differences found for orientation testing (Fig. 3a, all *p*_FDR_ *<* 0.05). These findings are consistent with the fact that the MoCA was designed to evaluate Alzheimer’s disease, which makes up a large portion of Cluster 2. Interestingly, the visuospatial subsection produced better separation in performance across multiple clusters. For this section, the Cluster 2 median was lower than the Cluster 1 median (Fig. 3a, *n* = 60, *M*_2−1_ = −2.00, *p*_FDR_ *<* 0.01) and than the Cluster 3 median (Fig. 3a, *n* = 58, *M*_2−3_ = − 2.30, *p*_FDR_ *<* 0.05). The Cluster 1 median and Cluster 3 median visuospatial scores were higher than the Cluster 4 median scores (Fig. 3a; *M*_1−4_ = 1.21, *n* = 89, *p*_FDR_ *<* 0.05; *M*_3−4_ = 1.50, *n* = 87, *p*_FDR_ *<* 0.05). These results suggest that testing of visuospatial cognition and executive function domains are more sensitive to variation in neuropathological disease subtype than other MoCA sections, which are primarily sensitive to Alzheimer’s disease.

**FIG. 3.**
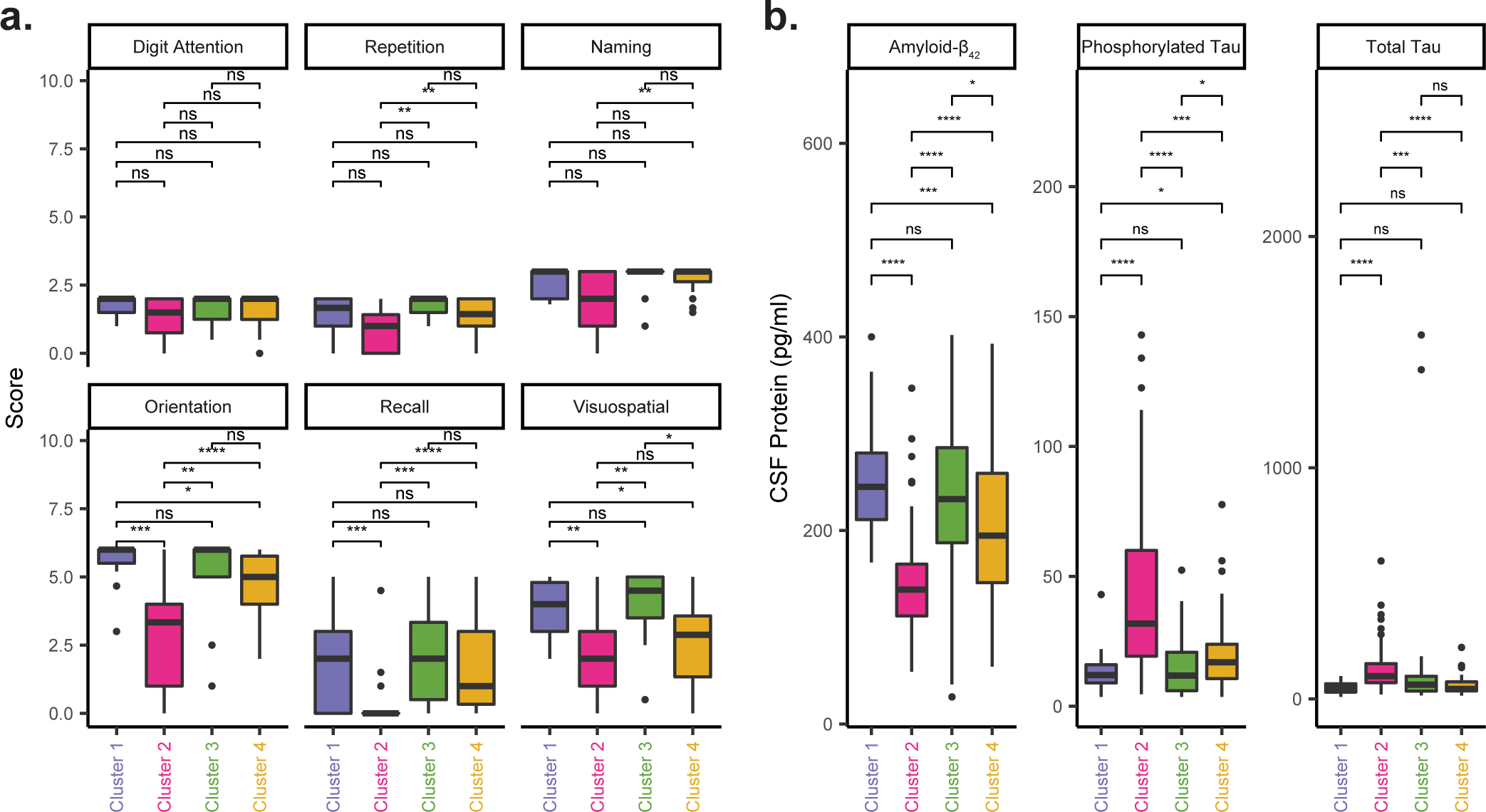
Cognitive deficits and CSF protein levels in disease clusters. *(a)* Pairwise intercluster comparisons of median Montreal Cognitive Assessment (MoCA) subscores^46^ using the Wilcoxon rank-sum test, FDR-corrected for multiple comparisons (*q <* 0.05) over all pairwise tests for 6 subscores. *(b)* Pairwise intercluster comparisons of median CSF protein levels for amyloid-*β*_1−42_, phosphorylated tau, and total tau using the Wilcoxon rank-sum test, FDR-corrected for multiple comparisons (*q <* 0.05) over all pairwise tests for all 3 proteins. These results were robust to the removal of the two subjects with total tau *>* 1000 pg/ml in Cluster 3 (Fig. S9). Plots were constructed using code from R package *ggpubr* ^47^. *ns, p*_FDR_ *>* 0.05. *, *p*_FDR_ *<* 0.05. **, *p*_FDR_ *<* 0.01. ***, *p*_FDR_ *<* 0.001. ****, *p*_FDR_ *<* 10^*−*6^. CSF, cerebrospinal fluid.

In addition to cognitive biomarkers of disease, we were also interested to know how our pathology-defined clusters separated patients with respect to the levels of proteins found in cerebrospinal fluid (CSF), a diagnostic test primarily used to identify Alzheimer’s disease^40,48–50^ with mixed success in FTLD^39^. Specifically, we assessed CSF amyloid-*β*_1−42_, phosphorylated tau, and total tau in an *n* = 268 subsample of patients with available data. Again, we used the Wilcoxon rank-sum test to evaluate the null hypothesis that the median sample difference between every unique pairwise combination of clusters is equal to 0 for each CSF protein, correcting for multiple comparisons over 6 unique tests for 3 CSF proteins by adjusting the false discovery rate (*q <* 0.05). We found statistically significant differences in median CSF amyloid-*β*_1−42_ between every pair of clusters except for Cluster 1 relative to Cluster 3 (Fig. 3b, all *p*_FDR_ *<* 0.05). Phosphorylated tau levels separated Cluster 2 from every other cluster (Fig. 3b; *M*_2−1_ = 19.00, *n* = 187, *p*_FDR_ *<* 10^*−*6^; *M*_2−3_ = 19.10, *n* = 189, *p*_FDR_ *<* 10^*−*6^; *M*_2−4_ = 13.85, *n* = 192, *p*_FDR_ *<* 0.001), as well as Cluster 4 from Clusters 1 and 3 (Fig. 3b; *M*_4−1_ = 4.34, *n* = 79, *p*_FDR_ *<* 0.05; *M*_4−1_ = 5.00, *n* = 81, *p*_FDR_ *<* 0.05). Total tau levels alone separated Cluster 2 from every other cluster (Fig. 3b; *M*_2−1_ = 52.00, *n* = 187, *p*_FDR_ *<* 10^*−*6^; *M*_2−3_ = 36.29, *n* = 189, *p*_FDR_ *<* 0.001; *M*_2−4_ = 48.00, *n* = 192, *p*_FDR_ *<* 10^*−*6^). These results are again consistent with the fact that the analysis of these particular CSF proteins was developed with Alzheimer’s disease pathophysiology in mind. It is particularly notable that amyloid-*β*_1−42_ and phosphorylated tau produced separation between nearly all 4 disease clusters, suggesting that certain ranges of CSF amyloid levels may be characteristic of particular underlying disease processes.

Finally, we hypothesized that by considering multiple forms of pathology in defining new disease categories, the patients within the resulting categories would exhibit phenotypic differences that transcend boundaries of existing disease labels. To test this hypothesis, we repeated our analysis of CSF protein levels while either excluding Alzheimer’s disease from Cluster 2 (Fig. S4a) or isolating patients with Alzheimer’s disease (Fig. S4b). Notably, this approach greatly reduced our statistical power, but we performed the analysis nevertheless to probe the phenotypic boundaries of our disease clusters. We defined Alzheimer’s disease as both Braak and CERAD scores *>* 1, requiring evidence of both neurofibrillary tangle and neuritic plaque burden, respectively. This diagnostic criterion is hereafter referred to as “Braak-CERAD Alzheimer’s disease.” In an *n* = 126 subsample with no patients in Cluster 2 meeting the Braak-CERAD Alzheimer’s disease definition, we found that Cluster 2 still demonstrated lower median CSF amyloid-*β*_1−42_ than Cluster 1 (Fig. S4a; *M*_2−1_ = −78.08, *n* = 45, *p*_FDR_ *<* 0.01) and than Cluster 3 (Fig. S4a, *M*_2−3_ = −71.82, *n* = 47, *p*_FDR_ *<* 0.05). Additionally, Cluster 2 still had a higher median CSF phosphorylated tau than Cluster 1 (Fig. S4a; *M*_2−1_ = 7.62, *n* = 45, *p*_FDR_ *<* 0.05). In an *n* = 159 subsample of patients exclusively meeting the Braak-CERAD Alzheimer’s disease definition, we did not find any statistically significant differences between Cluster 2 and Cluster 4 (the only two clusters with *>* 2 patients with Braak-CERAD Alzheimer’s disease and CSF protein data available). However, compared to Cluster 4, CSF amyloid-*β*_1−42_ was lower in Cluster 2 (Fig. S4b, *M*_2−4_ = −18.47, *n* = 159, *p*_FDR_ *>* 0.05) and total (Fig. S4b, *M*_2−4_ = 29.00, *n* = 159, *p*_FDR_ *>* 0.05) and phosphorylated tau (Fig. S4b, *M*_2−4_ = 10.46, *n* = 159, *p*_FDR_ *>* 0.05) were higher in Cluster 2, consistent with the pattern observed in Fig. 3b and Fig. S4a. In sum, these disease clusters may represent a new subtype of neurodegeneration, whose phenotypic boundaries cross Braak-CERAD Alzheimer’s disease. Overall, our findings highlight the transdiagnostic nature of the pathology-based disease clusters.

### Genotypic signatures of disease clusters

Genetic factors play an important role in determining risk for development of neurodegenerative disease. The *ϵ*4 allele at the apolipoprotein E (*APOE*) gene locus is a strong risk factor for Alzheimer’s disease, while the *ϵ*2 allele is thought to be protective^51^. Interestingly, the H1 haplotype at the gene locus encoding tau (*MAPT*) has been associated with progressive supranuclear palsy (PSP)^52^, Parkinson’s disease^53^, and Alzheimer’s disease^54^, three diseases with putatively distinct etiologies. We sought to understand how these genetic risk factors might be represented within our pathology-based disease clusters. Using a subsample of 1306 patients with genotyping at the *APOE* and *MAPT* loci, we measured the representation of alleles within each cluster (Fig. 4a-b). Next, using multiple logistic regression, we measured the odds of cluster membership given the presence of risk alleles.

**FIG. 4.**
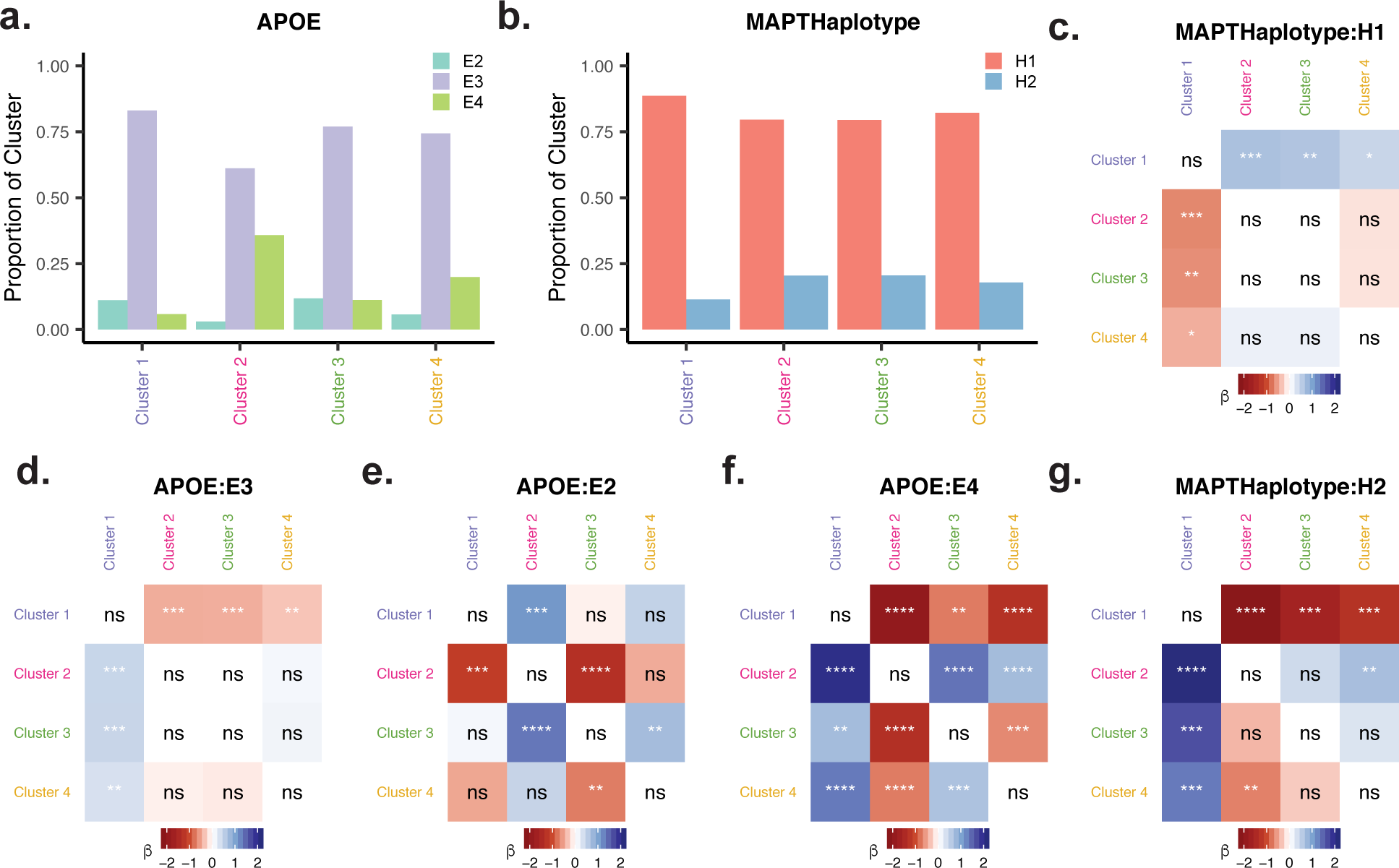
Prevalence of Alzheimer’s disease risk alleles differs across disease clusters. *(a-b)* Within each cluster, we calculated the proportion of each allele for *APOE* (panel *a*) and *MAPT* (panel *b*). *(c,g)* Matrix of logistic regression *β*-weights, whose *ij*-th element reflects the increase in log odds ratio for membership to cluster *i* relative to cluster *j* given the presence of *MAPT* -H2 (panel *g*) or *MAPT* -H1 (panel *c*). *(d-f)* Matrix of logistic regression *β*-weights, whose *ij*-th element reflects the increase in log odds ratio for membership to cluster *i* relative to cluster *j* given the presence of *APOE* -*ϵ*2 (panel *e*), *APOE* -*ϵ*3 (panel *d*), or *APOE* -*ϵ*4 (panel *f*). ns, *p*_FDR_ *>* 0.05. *, *p*_FDR_ *<* 0.05. **, *p*_FDR_ *<* 0.01. ***, *p*_FDR_ *<* 0.001. ****, *p*_FDR_ *<* 10^*−*6^.

First, we used this approach to investigate how *APOE* alleles were distributed across our disease clusters. We found that subjects carrying an *APOE* -*ϵ*4 allele had higher odds of belonging to Cluster 2 than to any other cluster (Fig. 4f, all *p*_FDR_ *<* 10^*−*6^; Cluster 1, *β* = 2.01, *df* = 669; Cluster 3, *β* = 1.38, *df* = 815; Cluster 4: *β* = 0.73, *df* = 803). Also, subjects carrying an *APOE* – *ϵ*2 allele had lower odds of belonging to Cluster 2 than to Cluster 1 or Cluster 3 (Fig. 4e; Cluster 1, *β* = −1.11, *df* = 669, *p*_FDR_ *<* 0.001; Cluster 3, *β* = −1.36, *df* = 815, *p*_FDR_ *<* 10^*−*6^). Importantly, when we carried out the same analysis in an *n* = 891 subsample with no Braak-CERAD Alzheimer’s disease patients in Cluster 2, the remaining patients in Cluster 2 were still more likely to carry an *E*4 allele relative to Cluster 1 and Cluster 3 (Fig. S5e; Cluster 1, *β* = 1.69, *df* = 248, *p*_FDR_ *<* 10^*−*6^; Cluster 3, *β* = 1.08, *df* = 394, *p*_FDR_ *<* 0.001, respectively), supporting the transdiagnostic nature of our disease clusters. These results suggest that *APOE* -*ϵ*4 may not carry risk for Alzheimer’s disease *per se*, but rather for a syndrome with alternative boundaries with respect to molecular and cellular pathology.

Next, we used the same regression-based approach to examine how *MAPT* alleles were distributed across our disease clusters. We found that the presence of two H2 alleles portended lower odds of Cluster 1 membership relative to all other clusters (Fig. 4g; Cluster 2, *β* = −2.14, *df* = 670, *p*_FDR_ *<* 10^*−*6^; Cluster 3, *β* = −1.67, *df* = 504, *p*_FDR_ *<* 0.001; Cluster 4, *β* = −1.33, *df* = 492, *p*_FDR_ *<* 0.001). Similarly, the odds of Cluster 1 membership were higher given the presence of an H1 allele relative to all other clusters (Fig. 4c; Cluster 2, *β* = 0.67, *df* = 670, *p*_FDR_ *<* 0.001; Cluster 3, *β* = 0.64, *df* = 504, *p*_FDR_ *<* 0.01; Cluster 4, *β* = 0.45, *df* = 492, *p*_FDR_ *<* 0.05). PSP is known to be associated with the *MAPT* H1 haplotype and was primarily found in Cluster 1. Therefore, we repeated the above analysis while excluding PSP patients from Cluster 1 to test whether the associations between Cluster 1 membership and *MAPT* haplotypes could be simply explained by the sequestration of PSP patients in Cluster 1. When excluding PSP from Cluster 1, the relationship between the odds of Cluster 1 membership and the presence of an H2 allele was still present (Fig. S6a; Cluster 2, *β* = −1.96, *df* = 597, *p*_FDR_ *<* 10^*−*6^; Cluster 3, *β* = −1.53, *df* = 431, *p*_FDR_ *<* 0.001; Cluster 4, *β* = −1.23, *df* = 419, *p*_FDR_ *<* 0.01), although the relationship between Cluster 1 odds and H1 allele presence was weakened (Fig. S6a, all *p*_FDR_ *>* 0.05; Cluster 2, *β* = 0.27, *df* = 597; Cluster 3, *β* = 0.25, *df* = 431; Cluster 4, *β* = 0.09, *df* = 419). Overall, these findings suggest that our pathology-based clusters automatically produced categories whose genotypic compositions cross boundaries drawn by existing disease labels.

### Multivariate classification of disease labels from *in vivo* biomarkers

Clinicians currently utilize CSF protein analysis and genotyping in diagnosing neurodegenerative disease^6,40,48^. However, heterogeneity within existing disease categories hinders the use of these tests to accurately infer a specific underlying histopathologic syndrome afflicting a patient with putative dementia. We were interested in quantifying the utility of *in vivo* biomarkers in predicting histopathologic disease labels from our clustering solution compared with labels from existing disease definitions. In 268 patients with available data, we trained a multiple logistic regression model to identify a single class of disease label out of the remaining heterogeneous group of patients using CSF amyloid-*β*_1−42_, phosphorylated tau, and total tau protein levels. In this way, we convert a multi-class prediction problem to a one-vs.-all scenario to increase the available sample size for model training. Additionally, compared to a pairwise comparison between two disease entities^49,50^, a one-vs.-all prediction is both more difficult and contributes more information if performed accurately, as it requires no clinical priors or narrowing of problem scope to a particular syndrome.

Interestingly, we found that the out-of-sample area under the receiver-operater characteristic curve (AUC) was very similar with the inclusion of genotype data (Fig. S7), suggesting that CSF protein levels and *APOE* /*MAPT* genotype explain common variance in disease status. Notably, however, disease labels can still be predicted from genotype alone with above-chance accuracy (Fig. S10). We achieved similar accuracy using random forest^55^, an algorithm capable of learning nonlinear relationships between classes and features (Fig. S8). Here, we present the out-of-sample performance of a multiple logistic regression model using only CSF protein levels as features. The model was able to identify a neuropathologic diagnosis of Alzheimer’s disease (defined as both Braak and CERAD scores *>* 1) with mean AUC = 0.94 (Fig. 5a, c). Performance in identifying neuropathologic diagnoses of Lewy body dementia (LBD) or FTLD-TDP was weaker yet still above chance, with mean AUC of 0.70 and 0.75, respectively (Fig. 5a, c). We did not attempt to predict diseases with fewer than 25 patients with CSF protein data. Collectively, these findings suggest that CSF protein can be used to distinguish patients with a particular traditional neuropathologic diagnosis from a group of patients with a heterogeneous mix of neurodegenerative diseases.

**FIG. 5.**
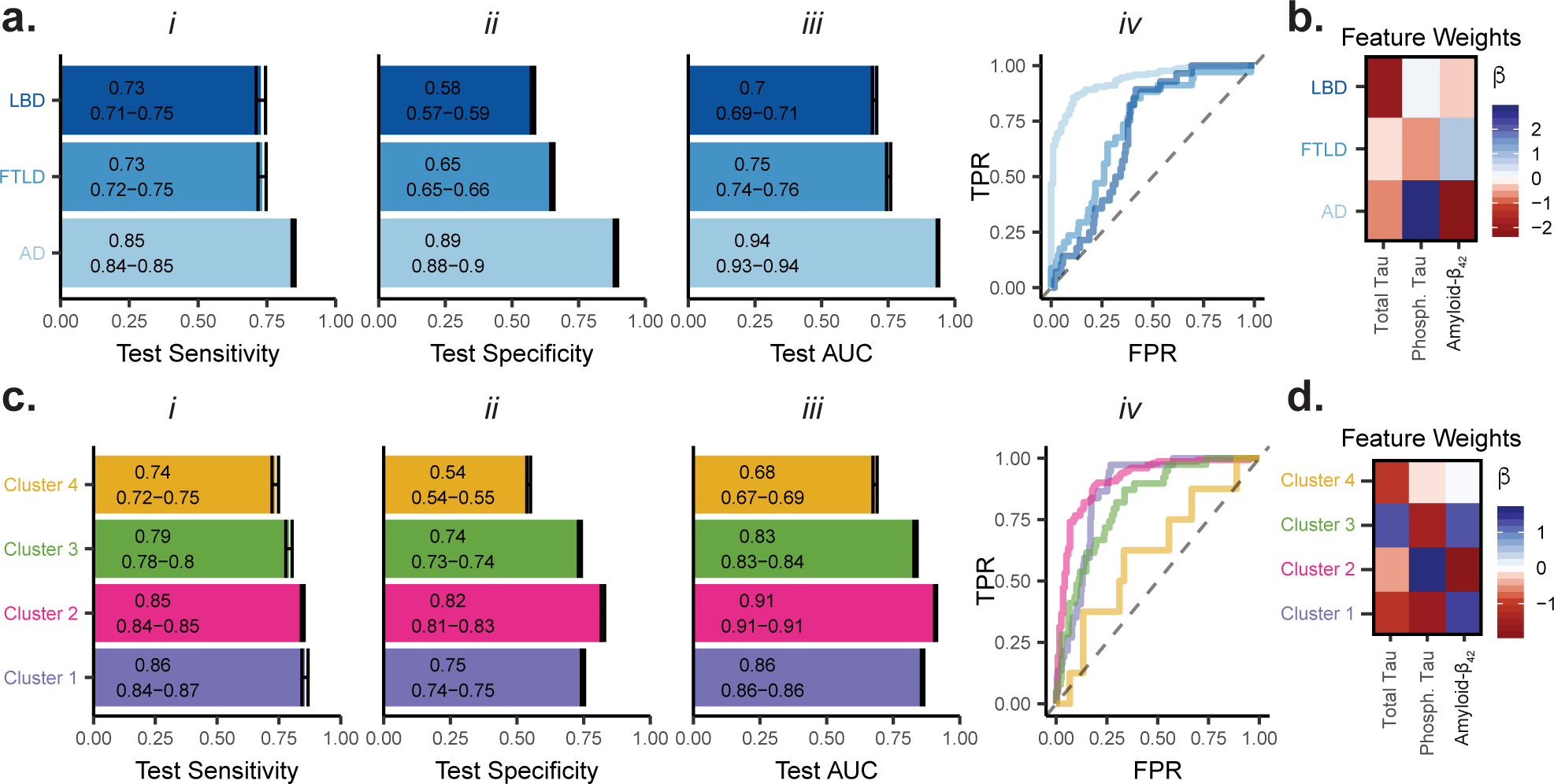
Identifying disease labels from initial testing of CSF protein. *(a, c)* Characteristics of prediction of existing diagnoses *(panel a)* or disease clusters *(panel c)* in held-out testing data using multiple logistic regression to predict disease labels from CSF protein levels. Sub-panels *i* and *ii* show the test-set sensitivity and specificity, respectively, using a threshold value of 0.5. Sub-panel *iii* shows the area under the curve (AUC) on the test-set, reflecting performance over a range of threshold values. Bar length represents mean performance, and error bars indicate 95% confidence intervals over 100 repetitions of *k*-fold cross-validation at *k* = 10. Sub-panel *iv* shows representative receiver-operator characteristic curves for test-set predictions of existing diagnoses *(panel a)* or disease clusters *(panel c). (b, d)* Mean standardized multiple logistic regression *β* across 100 repetitions of *k*-fold cross-validation at *k* = 10 in the prediction task for existing disease labels *(panel e)* or disease clusters *(panel f)*. The *β* weights can be interpreted as the increase in log-odds ratio for a one standard deviation increase in the value of the predictor. TPR, true positive rate. FPR, false positive rate. Total Tau, total CSF tau protein. Phosph. Tau, total CSF phosphorylated tau. Amyloid-*β*_1−42_, total CSF amyloid-*β*_1−42_.

After demonstrating that we could predict existing disease labels with high accuracy from CSF protein levels, we next asked whether the disease cluster we identified could improve our ability to infer histopathologic syndromes from clinical data. With the same group of patients, we carried out the same identification procedure using disease cluster membership as class labels instead of existing disease definitions. We achieved mean AUCs of 0.68, 0.83, 0.91, and 0.86 for each cluster, respectively (Fig. 5c-d). Using the data-driven disease clusters, we identified Cluster 3 with higher AUC by 0.08 than FTLD-TDP as defined under the traditional system of disease labels. We compare Cluster 3 to FTLD-TDP because 79.1% of FTLD-TDP patients were grouped into Cluster 3. Additionally, we could identify Cluster 1 membership (tauopathies, Fig. 2e) with AUC = 0.86, whereas no individual disease with meaningful representation in Cluster 1 had enough available patients for classification. These findings suggest that the disease cluster labels were resolved by CSF protein levels at least as well as traditional definitions of neurodegenerative disease. The disease cluster labels also allow for prediction of tauopathy family membership by automatically grouping together related diseases.

Next, we aimed to explore how the transdiagnostic nature of our clusters interacted with our ability to classify patients using CSF protein. Notably, the AUC for Cluster 2 was similar to that for Alzheimer’s disease, consistent with Cluster 2’s primary composition of Alzheimer’s disease patients (Fig. 2e). However, Cluster 2 also includes a heterogeneous mix of patients without a diagnosis of Braak-CERAD Alzheimer’s disease. Therefore, despite displaying slightly lower (0.03) mean AUC than the identification of Braak-CERAD Alzheimer’s disease, Cluster 2 identification (AUC = 0.91) is impressive in light of Cluster 2’s heterogeneous composition. Furthermore, when we exclude patients with Braak-CERAD Alzheimer’s disease from Cluster 2, we could still predict the heterogeneous subsample of Cluster 2 with accuracy above chance (Fig. S11, AUC = 0.74) despite a greatly reduced sample size. Overall, these findings indicate that we can infer membership to 4 neurodegenerative disease clusters, each characterized by a particular pathogenic protein process, with high accuracy solely from CSF protein.

Finally, in order to understand how each model classified certain patients into disease clusters compared with existing diagnoses, we examined the feature weights from our multiple logistic regression model. Feature weights for Cluster 1 and Cluster 2 were similar to those for Alzheimer’s disease and FTLD, respectively. This finding suggests that the ability to accurately identify patients falling into alternative histopathologic boundaries relies on slight readjustments to the weight on each CSF feature, without drastically altering the ratios between feature weights.

## DISCUSSION

Neurodegenerative diseases are typically defined by the increased burden of one or two pathogenic protein species. However, it is the rule rather than the exception that individual patients meet criteria for multiple diseases and exhibit several pathogenic protein aggregates. In the present work, we analyzed a large post-mortem sample of patients with a diverse representation of neurodegenerative diseases. Using a disease-blind approach, we assigned each patient to a new disease cluster by simultaneously accounting for the levels of 6 key aggregated protein species and 3 histological features across 18 regions. The resulting clusters, defined only from pathology data, differed in terms of cognition, genetics, and CSF protein levels. Finally, we trained statistical models that were able to classify cluster membership with above-chance accuracy from *in vivo* measurements. This work represents an advance in our understanding of the clinical and neuropathologic heterogeneity among neurodegenerative diseases. Furthermore, our methods and approach provide a general framework that could be applied to various clinical populations outside of patients with neurodegeneration.

### Accounting for copathology produces transdiagnostic categories of neurodegenerative disease

Existing definitions of neurodegenerative disease typically require the presence of one or two pathological proteins and reflect decades of clinical and scientific consensus^4,6,7^. Faced with the reality of co-occurring pathology, this system will assign multiple disease labels to a single patient, making it difficult to identify clinical phenotypes and genetic variants associated with neurodegenerative processes. In the present work, we define 4 non-overlapping disease categories using an unsupervised approach that simultaneously considers 162 pathological features (Fig. 2). These categories appear to be primarily driven by global levels of 4 proteins (Fig. S3; Cluster 1: tau, Cluster 2: amyloid-*β*, Cluster 3: TDP-43, and Cluster 4: *α*-synuclein), rather than their regional distributions.

We found that our clustering approach groups together existing disease entities caused by each pathological protein, while simultaneously parsing heterogeneity both within and between diseases. Patients with Alzheimer’s disease were found in all 4 clusters, though primarily in Cluster 2. However, nearly all patients with Alzheimer’s disease in Cluster 4 had a secondary diagnosis of LBD, while most Alzheimer’s patients in Cluster 1 and 3 had no additional diagnoses. Additionally, a small cohort of Alzheimer’s disease patients within Cluster 2 also carried a secondary diagnosis of LBD, though most harbored no additional diagnoses. The bifurcation of patients with concomitant amyloid-*β*, tau, and *α*-synuclein pathology into Cluster 2 and Cluster 4 could reflect the existence of multiple strains of *α*-synuclein^56^, which may differentially interact with tau^57^.

Despite being defined solely from pathology data, our clusters separated patients phenotypically and genotypically. We found that mean CSF amyloid-*β* and *APOE* -*ϵ*4 allele representation differed between almost every pair of clusters (Fig. 2 and 3). These differences are inherently transdiagnostic due to the cluster composition, but we confirmed that differences involving Cluster 2 were not trivially driven by the large Alzheimer’s disease component. Indeed, Cluster 2 members who did not meet criteria for Alzheimer’s disease also exhibited increased *APOE* -*ϵ*4 prevalence (Fig. S5) and reduced CSF amyloid-*β* relative to other clusters (Fig. S4). This separation of Cluster 2 is likely in part achieved by the relative exclusion of Alzheimer’s disease patients from other clusters, as well as by the inclusion of patients in Cluster 2 who did not meet criteria for Alzheimer’s disease but may bear pathological similarity with respect to cellular features or sub-diagnostic levels of amyloid-*β* and tau. Overall, these phenotypic and genotypic differences support a clinical and biological relevance for our copathology-driven disease clusters. Our results suggest that low CSF amyloid-*β* and the *APOE* -*ϵ*4 allele, markers traditionally associated with Alzheimer’s disease, might in reality reflect a pathological syndrome with slightly different histopathological boundaries than Alzheimer’s disease.

### Statistical models expand the utility of CSF protein analysis

An ideal route towards targeted therapies generally involves the identification of a sufficiently homogeneous clinical population whose biological characteristics motivate the use of a targeted treatment. In the case of neurodegenerative disease, phenotypic and genotypic heterogeneity, along with discordance between clinical diagnoses and gold-standard neuropathological diagnoses^39^, limit a clinician’s ability to map biomarkers to specific neurodegenerative processes. To address this problem, we trained generalized linear models to predict histopathologic disease class membership based on CSF protein levels and genotype at the *APOE* and *MAPT* loci. Notably, for all predictive modeling, we included all available patients regardless of diagnosis, used the chronologically earliest available CSF sample, and reported model performance on a distribution of held-out samples in order to mirror a realistic clinical scenario. We were able to identify Braak-CERAD Alzheimer’s disease (AUC = 0.94) and Cluster 2 membership (AUC = 0.91) from a heterogeneous clinical population of patients with neurodegenerative disease using CSF protein levels (Fig. 5a,c). The high accuracy prediction of Cluster 2 membership is impressive in light of its data-driven construction and heterogeneous composition of overlapping traditional diagnoses. Predicting this alternative category only requires slight readjustments to the weights on each CSF feature, without perturbing the relative directionality of the weights (Fig. 5e-f). Additionally, while individual tauopathies were too sparsely represented to train predictive models that could be robustly tested out-of-sample, the disease cluster system allowed us to accrately identify membership to a family of tauopathies (AUC = 0.86). Finally, we demonstrated above chance accuracy in disease class identification based solely on genotype at two loci.

All in all, these findings suggest an untapped utility in genotyping and CSF protein analysis for identifying subgroups of patients within flexibly defined histopathologic boundaries, using simple statistical models to generate predictions. Such models may be valuable for designing and repurposing pharmaceuticals, immunotherapies, or combination therapies^58^ by allowing their efficacy to be compared against probabilistic estimates of membership to a particular histopathologic disease class. Moreover, predictions based on genotype would be stable throughout an individual’s life span, allowing for prospective evaluation of treatments designed to intervene early in the disease course.

### Limitations

We acknowledge a number of limitations in the present study. First, the semiquantitative assessment of the degree of each pathological feature precludes the use of statistical methods relying on interval or continuous data and is subject to issues of inter-rater reliability. More granularity in the levels of pathology might be obtained through quantitative, automated image analysis^59,60^. Continuous pathology data would lend itself well to principal component analysis, which might identify dimensions of covarying neuropathological features. The loadings on these dimensions could then be correlated with phenotype in a continuous fashion, rather than generating a hard partition and performing comparisons of means, as in the present work. However, such approaches for quantitative mapping are still being validated and the large sample size of manually labeled data vastly outweighs the benefit of using smaller amounts of automatically labeled data. The fact that we are able to generate meaningful disease categories consistent with prior work argues that the use of ordinal data is not a drastic limitation, and may inspire others to use similar approaches on datasets that have remained incompletely explored due to their discrete nature.

Another caveat is that the cluster definitions depend on the sample composition. However, application of the same approach to samples with different compositions might yield independently interesting results by relative subtyping. Nevertheless, a new observation (from any new patient) could be assigned to one of the clusters analyzed in this paper by simply taking the maximum Spearman correlation with each cluster centroid.

Finally, the availability of phenotypic measurements (MoCA and CSF protein analysis) are biased by clinical decision making. One can imagine that lumbar puncture followed by CSF protein analysis and detailed cognitive phenotyping were not routinely performed on patients primarily exhibiting motor symptoms. However, our sample size was large enough that we were able to validate our model in a distribution of held-out samples, unlike prior applications of CSF-based predictive models which report training set accuracy in smaller samples with well-circumscribed disease boundaries^49,50^. This validation procedure increases confidence in the external validity of our model. Nevertheless, the true model performance “in the wild” cannot be accurately estimated without external validation in an independent sample.

### Future directions

The utility of the statistical models in the present work are limited by sample size, but also in the availability of relevant features. Incorporation of quantitative data on clinical symptomatology, along with more complete genomic data, would likely also enhance the predictive value of these models. Several of the disorders studied in the present work can be identified clinically, and are associated with multiple genetic variants^6,61,62^. Clinical symptomatology and genotype can be obtained noninvasively, and might be applied more easily as well as explain additional variance in disease. In particular, a model based purely on genotype could theoretically provide an estimate of disease risk at birth, which would allow for the development of preventative therapies targeting early, preclinical disease.

Furthermore, the general approach we utilized in the present work is neither specific nor limited to neurodegenerative disease. Unsupervised learning can be applied to similarity matrices based upon any set of features pertinent to a disease with overlapping pathophysiologic modes, as in multifactorial disorders such as epilepsy, vascular disease, or cancer. Crucial to this endeavor is the compilation of large multimodal and multisite datasets that capture a broad range of diagnoses, phenotypes, and genotypes. In addition to the potential clinical utility of biomarker-based forecasting of histopathological syndromes, the present work may serve as a model for the use of unsupervised methods to identify data-driven, transdiagnostic disease subtypes in any field.

## METHODS

### Sample construction

All data were obtained through the Integrated Neurodegenerative Disease Database (INDD)^41,63^, hosted by the Center for Neurodegenerative Disease Research at the Hospital of the University of Pennsylvania. A team of expert neuropathologists assessed the extent of 6 molecular pathological features (amyloid-*β*, neuritic plaques, tau, *α*-synuclein, TDP-43, and ubiquitin) and 3 cellular pathological features (angiopathy, neuron loss, and gliosis) for 18 brain regions during autopsy for 1757 patients as previously described^41,63^. Semiquantitative pathology scores (0, Rare, 1+, 2+, 3+) were converted to integers from 1 to 5 and treated as ordinal data except when computing mean regional pathology scores to visualize cluster centroids. Individuals with missing data for *>* 75% of pathological features were excluded from further analysis, and all available pathological features were included in all analysis (Fig. S1a-b, see Supplementary Information for discussion).

Expert neuropathologists assigned up to 4 histopathologic diagnoses to each patient according to well-accepted disease definitions codified in the neuropathology literature; namely, Alzheimer’s disease^4^, amyotrophic lateral sclerosis (ALS)^6^, argyrophilic grain disease^6^, cerbral amyloid angiopathy (CAA)^4^, cerebrovascular disease (CVD)^4^, frontotemporal lobar dementia with TDP-43 inclusions (FTLD-TDP)^6^, frontotemporal lobar dementia without TDP-43 inclusions (FTLD-Other)^6^, dementia with Lewy bodies (LBD)^64^, Parkinson’s disease with and without dementia^64^, multiple system atrophy^7^, primary age-related tauopathy (PART)^65^, Pick’s disease^6^, and progressive supranuclear palsy (PSP)^6^. We combined LBD and Parkinson’s disease with dementia for simplicity. All patients were also given a clinical diagnosis by their physician prior to autopsy.

### Unsupervised clustering of patients by molecular pathology

Traditional definitions of neurodegenerative diseases typically only account for 1-2 species of protein aggregate. Here, we sought to group patients into clusters using all available pathological features in an unsupervised manner^66–68^. Thus, we constructed a matrix **W** whose elements **W**_*ij*_ equal the Spearman correlation between vectors of all pairwise complete observations of 152 regional pathology scores for patient *i* and patient *j*. Before computing the Spearman correlation, we demeaned each feature to control for differences in the relative meaning of each score (i.e., a 1+ for amygdala tau may not correspond to the same relative burden as a 1+ for angular gyrus neuritic plaques). Next, we performed 1000 iterations of modularity maximization^69^ on the matrix **W**, using a definition of modularity designed for networks with both positive and negative weights^70,71^. This definition of modularity considers the positive and negative weights in **W**_*ij*_ separately, such that the absolute values of the positive and negative weights in **W**_*ij*_ are denoted as 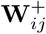 and 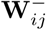, respectively. The total weight of the network, 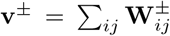, is the sum of all positive or negative weights. The strength of node *i*, 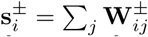, is the sum of all positive or negative weights at node *i*. Then, modularity is defined as

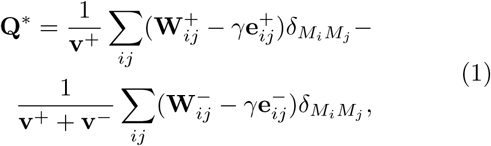

where the expected positive or negative connection weights within modules are **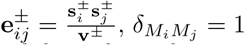** when *i* and *j* are in the same module and 0 otherwise, and *γ* is a resolution parameter^72^. This approach^71^ uses a Louvain-like^73^ locally greedy algorithm to maximize **Q**^*\**^ through the selection of a partition *M*. In this instantiation of modularity maximization, larger values of *γ* result in the identification of many small clusters, while smaller values of *γ* result in the identification of a few larger clusters. We selected the *γ* value that gave us the highest partition stability, reflected by the mean *z*-scored Rand index^74^ between all partitions generated at a particular value of *γ*^75^. Partition stability peaked between γ values of 1.5 and 1.8. Accordingly, we present results for *γ* = 1.7 in the main text (see Supporting Information and Fig. S2 for further discussion).

### Group comparisons of phenotypic measurements

We used non-parametric testing to compare phenotypes between disease clusters. Using pairwise Wilcoxon rank-sum tests, we compared mean scores in 6 subdomains of the Montreal Cognitive Assessment^46^ (MoCA) between all unique combinations of clusters. All *p*-values were FDR-corrected (*q <* 0.05) over all comparisons for all 6 subdomains. The abstraction score was excluded due to missing data. We also used pairwise Wilcoxon rank-sum tests to compare CSF protein levels between all unique combinations of clusters. For patients that underwent multiple CSF studies, we only considered their earliest measurement in order to control for disease progression as much as possible. All *p*-values were FDR-corrected (*q <* 0.05) over all comparisons for the 3 CSF proteins that we assessed.

### Logistic regression of cluster membership on allele counts

Assuming a multiplicative genomics model^76^, we used multiple logistic regression to perform pairwise comparisons of allele distributions between disease cluster membership as binary phenotypes^77^. First, for each pair of clusters *i* = 1,.., *k* and *j* = 1,.., *k*, we constructed a *N*_*ij*_ *×* 1 binary outcome vector *Y*_*ij*_, where *N*_*ij*_ is the number of patients belonging to cluster *i* or cluster *j*. The elements of *Y*_*ij*_ indicate whether a patient belongs to cluster *i* or cluster *j*. A value of 1 indicates membership to cluster *i* and a value of 0 indicates membership to cluster *j*. Next, we construct a corresponding *N*_*ij*_*× n*_*g*_ allele table *A*, where *n*_*g*_ is the number of non-wild type alleles for a given gene *g*. The table *A* consists of *n*_*g*_ column vectors, *A*_*a*_, where *a* = 1,.., *n*_*g*_, and the elements of each column vector *A*_*a*_ indicate how many copies of allele *a* of gene *g* are present in each of the *N*_*ij*_ patients belonging to cluster *i* or cluster *j*. Next, we assumed additive genetic effects and used logistic regression^77^ to fit the following model to predict *Y*_*ij*_ from an allele table *A*_*g*_ for each gene *g*:

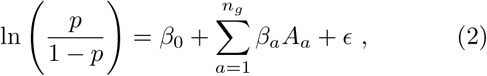

where *p* is the probability that a patient belongs to cluster *i, β*_0_ and *β*_*a*_ are parameters obtained by fitting the model, ln is the natural logarithmic function, and *E* is an error term. We fit this model for all unique pairwise comparisons of clusters, such that *ij* correspond to _*k*_*C*_2_. In this model, *β*_0_ is interpreted as the log odds that a patient belongs to cluster *i* given that the patient has two wild-type alleles. The parameter *β*_*a*_ is the change in log odds that a patient belongs to cluster *i* given the presence of 1 copy of allele *a*. We adjusted *p*-values over all comparisons for all genes and all alleles to control the false discovery rate at *q <* 0.05.

### Supervised learning of disease labels

In order to demonstrate a practical clinical use for multivariate models of biomarkers, we used data available *in vivo* to predict whether a patient met criteria for a specific neurodegenerative disease or was classified into a particular data-driven disease cluster. We converted this multi-class prediction scenario into a one-versus-all binary prediction scenario, in which we attempted to predict a *N ×* 1 binary disease label vector *D*_*i*_, whose elements equal 1 for patients positive for the *i*^th^ disease label and equal 0 for patients negative for the *i*^th^ disease label. Here, the disease labels were Alzheimer’s disease, LBD, FTLD-TDP, and the four clusters, indexed by *i* = 1,.., 7, and *N* is the number of patients with available data, either CSF protein levels, allele counts, or both. We used multiple logistic regression to predict ln *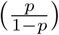*, where *p* is the probability that a patient is positive for the *i*^th^ disease label, from a feature matrix **X**, whose dimensions are *N × q*, where *N* is the number of patients with available data and *q* is the number of features. Depending on the analysis, **X** contained allele counts for non-wild-type alleles of 2 genes (*N* = 1312, *q* = 3), levels of 3 CSF proteins at initial evaluation (*N* = 268, *q* = 3), or both allele count and CSF protein levels (*N* = 262, *q* = 6).

When evaluating the performance of a model in predicting the *i*^th^ disease label, we wanted to ensure that our results were robust across multiple patient samples and consistent in a held-out sample of patients that were not involved in training the model. Therefore, we used 100 repetitions of *k*-fold cross validation^78^ to obtain estimates and confidence intervals for the out-of-sample performance of each model. For each repetition, we randomly split **X** and *D*_*i*_ along their rows into *k* = 10 subsets **X**_*j*_ and *D*_*j*_ for *j* = 1,.., *k*. When the *j*^th^ subset was used as the held-out testing dataset, **X**_*j*_ and *D*_*j*_ were the independent and dependent variables, respectively. The remaining *k −* 1 subsets were compiled into a training dataset, whose independent and dependent variables we refer to as **X**_r_ and *D*_r_. In order to aid the training process, we ensured class balance in **X**_r_ and *D*_r_ by randomly under-sampling their rows^79,80^, such that 50% of patients in **X**_r_ and *D*_r_ were positive for disease label *i* prior to training. Using multiple logistic regression, we fit the following equation with the training data:

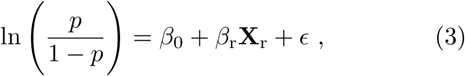

where *p* is a vector of probabilities that each patient is positive for the *i*^th^ disease label, *β*_0_ is an intercept vector, *β*_r_ is a vector of feature weights obtained through model fitting, ln is the natural logarithmic function, and *E* is a vector of random error terms. Next, we used the intercept *β*_0_ and feature weights *β*_r_ obtained by fitting our model to the training data to evaluate the out-of-sample performance of our model. We used the held-out testing data **X**_*j*_ to compute *D*_*p*_ = *β*_0_ + *β*_r_**X**_**j**_, where *D*_*p*_ is a vector of predicted log odds that each patient is positive for the *i*^th^ disease label. We converted *D*_*p*_ from a vector of log odds to a vector of probabilities and evaluated the performance of *D*_*p*_ in predicting *D*_*j*_. We repeated this procedure for *j* = 1,.., *k* until each subset *D*_*j*_ and **X**_*j*_ was used as the held-out data exactly one time. The entire *k*-fold cross-validation procedure was repeated 100 times for the *i*^th^ disease label to generate a distribution of test-set performance metrics (sensitivity, specificity, area under the curve). Reported sensitivity, specificity, and accuracy were obtained using a classification threshold value of 0.5 for *D*_*p*_, although prediction characteristics for a range of threshold values from 0 to 1 can be found in the receiver-operator characteristic curves.

## ACKNOWLEDGEMENTS

D.S.B. and E.J.C. acknowledge support from the John D. and Catherine T. MacArthur Foundation, the Alfred P. Sloan Foundation, the ISI Foundation, the Paul Allen Foundation, the Army Research Laboratory (W911NF-10-2-0022), the Army Research Office (Bassett-W911NF-14-1-0679, Grafton-W911NF-16-1-0474, DCIST-W911NF-17-2-0181), the Office of Naval Research, the National Institute of Mental Health (2-R01-DC-009209-11, R01 – MH112847, R01-MH107235, R21-M MH-106799), the National Institute of Child Health and Human Development (1R01HD086888-01), National Institute of Neurological Disorders and Stroke (R01 NS099348), and the National Science Foundation (BCS-1441502, BCS-1430087, NSF PHY-1554488 and BCS-1631550). E.J.C. also acknowledges support from the National Institute of Mental Health (F30 MH118871-01). J.Q.T. and V.M.Y.L. thank members of the Center for Neurodegenerative Disease Research who have contributed to the studies reviewed here. J.Q.T. and V.M.Y.L. also thank the patients and their families for brain donation. J.Q.T. and V.M.Y.L. acknowledge funding support from AG10124, AG17586, and the Woods Foundation. The content is solely the responsibility of the authors and does not necessarily represent the official views of any of the funding agencies.

## SUPPORTING INFORMATION

### Treatment of missing data

We examined the representation of missing values within the data to identify a rule by which we could choose subjects so as to maximize our sample size while still being able to compute a complete similarity matrix (Fig. 2a-c). When the pairwise available features between two patients lack variance, a correlation cannot be computed. This issue arises when a patient has uniform pathology and a small number of available features. Missing data was especially prevalent with features involving the dentate gyrus, occipital cortex, and ubiquitin, whose sampling was discontinued as a result of updated autopsy protocols. Notably, including additional features only increases the likelihood that a correlation can be computed and therefore we did not exclude any regions.

When we set a threshold for the maximum allowable percentage of features with missing data for each subject, we noticed that the number of available subjects did not begin to decrease substantially until a threshold value of 75% (Fig. S1a). Accordingly, we selected 75% as our threshold value such that no subject has missing data for *>*75% of the available features. We generated the same plot for thresholding the maximum allowable percentage of subjects with missing data for each feature, and we noticed that the number of available features also did not begin to decrease substantially until a threshold value of 75% (Fig. S1b). However, for reasons discussed above, we did not exclude any features.

### Optimization and validation of clustering procedure

In order to group patients into disease categories that simultaneously account for all measured forms of molecular and cellular pathology, we employed a graph-based clustering approach. First, we performed 1000 iterations of modularity maximization using a Louvain-like algorithm^71^ for *γ* values ranging from 0 to 3 in increments of 0.1. We computed the mean *z*-scored Rand index^74^ between all unique pairs of partitions at each *γ* value as a measure of partition stability at each resolution. We sought to choose the resolution with the highest mean similarity, because it can be more reliably and consistently generated. The highest mean similarity was reached at *γ* = 0.2, but this value appeared to be an unstable peak (Fig. S2a). However, we found a locally maximal plateau in mean similarity values between *γ* = 1.5 and *γ* = −1.8 (Fig. S2a). Accordingly, we proceeded with *γ* = 1.7. By examining a plot of the number of patients in each cluster as a function of *γ*, one can see that the partition at *γ* = 1.7 was generated for *γ* = 1.3 − 1.8, suggesting that the partition used in the main text does not depend on a narrow range of *γ* values.

Next, we demonstrated that our clustering approach was robust to outliers by generating 1000 *n* = 1111 sub-samples (80%) of our *n* = 1389 sample, demeaning the subsample, computing a Spearman correlation similarity matrix, and performing 50 iterations of modularity maximization at *γ* = 1.7 on that matrix followed by selection of the highest mean similarity partition. This procedure yielded 1000 subsampled partitions. We computed centroids for each partition and matched them to the most similar centroids from the full sample only if a set of unique matches could be made. Next, we plotted the distribution of correlations between the subsampled centroids and full-sample centroids (Fig. S2b), which showed that *>* 75% of the generated centroids had a Pearson corelation of near *r* = 1 with their respective best matched pair. These results suggest that the partition generated in the full sample and studied in the main text of the paper is robust to outliers, because it can be regenerated with subsets of the original data set.

### Disease clusters separate phenotypes and genotypes independent of existing disease labels

In order to probe the transdiagnostic nature of our disease clusters, we repeated the analyses in Fig. 3b and Fig. 4 while excluding or isolating certain existing disease entities. In this way, we examine how our disease cluster system parses heterogeneity both across and within existing disease boundaries.

First we compared CSF protein levels in an *n* = 126 subsample of patients from Cluster 2 after excluding patients with Braak-CERAD Alzheimer’s disease. We found that Cluster 2 still exhibited lower CSF amyloid-*β*_1−42_ than Cluster 1 (Fig. S4a, *p*_FDR_ *<* 0.01) and Cluster 3 (Fig. S4a, *p*_FDR_ *<* 0.05). Additionally, Cluster 2 had higher CSF phosphorylated tau than Cluster 1 (Fig. S4a, *p*_FDR_ *<* 0.05). These results suggest that non-Alzheimer’s disease Cluster 2 members are phenotypically similar to Alzheimer’s disease relative to other clusters.

When we isolated only individuals with Braak-CERAD Alzheimer’s disease, we had sufficient sample size (*n >* 2) to compare CSF protein levels between Cluster 2 and Cluster 4. While we did not find any statistically significant differences between Braak-CERAD Alzheimer’s disease in Cluster 2 versus Cluster 4, Cluster 2 exhibited a prototypical Alzheimer’s disease CSF profile, that is, low CSF amyloid-*β*_1−42_ with high CSF total tau and phosphorylated tau (Fig. S4b).

Next, we performed a similar analysis with respect to genotype. Specifically, we excluded Braak-CERAD Alzheimer’s disease from Cluster 2, yielding an *n* = 887 sample, and quantified the distribution of *APOE* and *MAPT* alleles across each cluster (Fig. S5a-b). Using pairwise multiple logistic regression, we estimated the increase in log odds ratio of cluster membership given the presence of *APOE* alleles (Fig. S5c-e). We found that the presence of an *APOE* -*ϵ*4 allele increased the odds of Cluster 2 membership relative to Cluster 1 (Fig. S5e, *p*_FDR_ *<* 10^*−*6^) and Cluster 3 (Fig. S5e, *p*_FDR_ *<* 0.001). These findings suggest that even Cluster 2 members with no diagnosis of Braak-CERAD Alzheimer’s disease were more likely to possess the *APOE ϵ*4 allele relative to other disease categories.

**FIG. S1.**
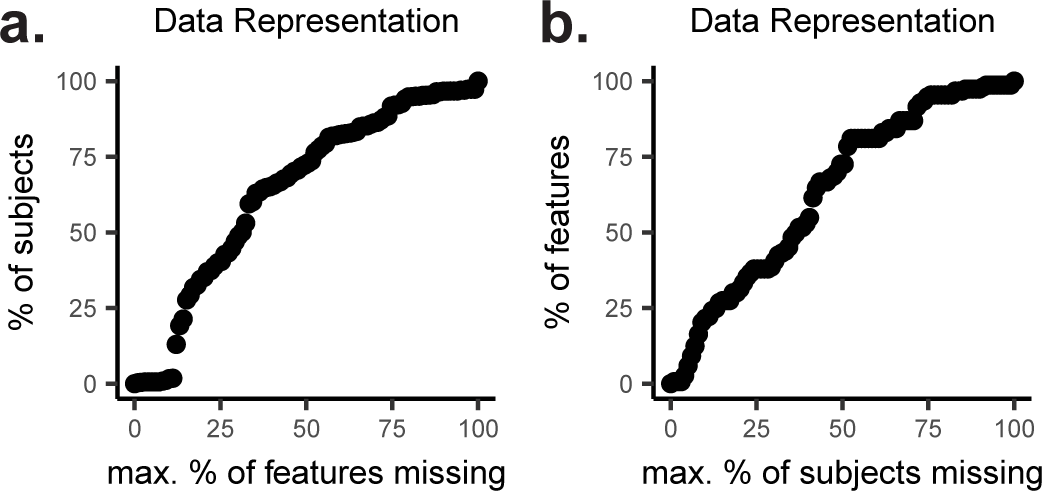
Missing data. *(a)* The *y*-axis shows the percentage of subjects available under different values on the *x*-axis for the maximum allowable percentage of missing features. *(b)* The *y*-axis shows the percentage of features available under different values on the *x*-axis for the maximum allowable percentage of missing subjects.

**FIG. S2.**
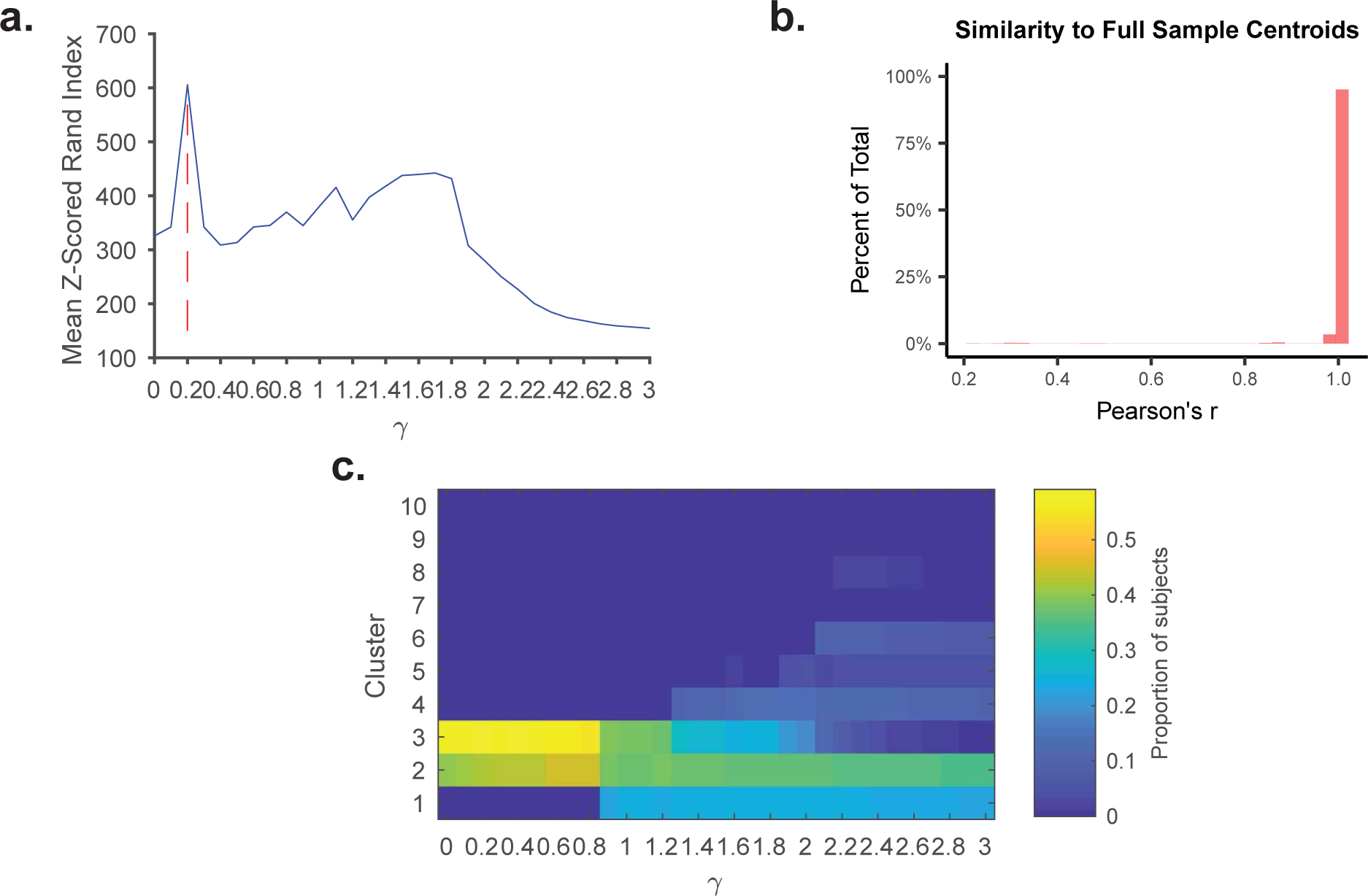
Choosing *γ* for modularity maximization. *(a)* Mean *z*-scored Rand Index between all unique combinations of *n* = 1000 partitions generated by applying modularity maximization for a range of *γ* values to the matrix of Spearman correlations in Fig. 2a. Blue line represents mean *z*-scored Rand index and red dashed line points to *γ* value with maximum mean *z*-scored Rand index. *(b)* Histogram of Pearson correlation values between 3986 centroids generated by carrying out the clustering procedure on 1000 80% subsamples of the *n* = 1389 sample and the original centroids from the full sample.

We also analyzed the extent to which progressive supranuclear palsy (PSP) was driving the genotypic associations involving Cluster 1. To answer this question, we excluded patients with PSP from Cluster 1, generating an *n* = 1239 sample, and then we used multiple logistic regression to compare the representation of *MAPT* alleles across clusters. We found that the log odds ratio of Cluster 1 membership was reduced given the presence of two H2 alleles at the H2 locus relative to Cluster 2 (Fig. S6a, *p*_FDR_ *<* 10^*−*^6), Cluster 3 (Fig. S6a, *p*_FDR_ *<* 0.001), and Cluster 4 (Fig. S6a, *p*_FDR_ *<* 0.01). We did not find statistically significant regression weights for H1 alleles as a predictor of Cluster 1 membership relative to any cluster, though the estimates were all greater than 0 (Fig. S6b, *p*_FDR_ *>* 0.05).

**FIG. S3.**
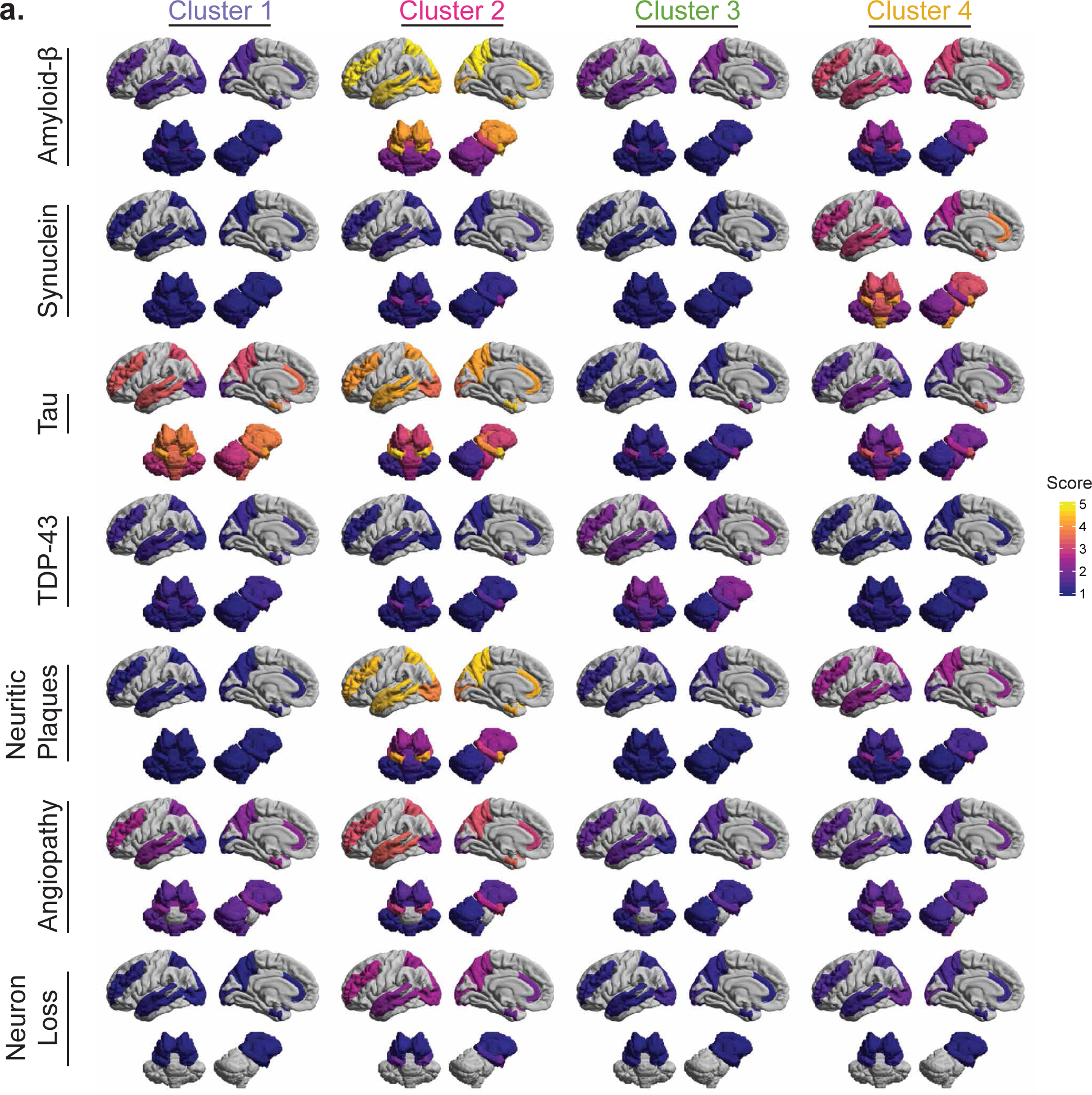
Spatial maps of representative disease cluster members. *(a)* We plotted mean regional pathology scores onto a surface rendering of a standard structural image^44^ using an anatomical parcellation^45^ to map autopsy sample sites^41^. Color-scaled mean pathology scores for each region and each pathological feature are shown for each disease cluster. In general, pathology was quantified for only one hemisphere, chosen at random, and therefore only one cortical hemisphere is shown and subcortical structures are plotted symmetrically for ease of visualization. Pathology scores from the substantia nigra and locus coeruleus were included in the data-driven clustering procedure, but those areas are not visualized here. Additionally, the dentate gyrus and CA1/subiculum were treated separately in the data-driven clustering procedure, but here are plotted as an average on the entire hippocampus. See Fig. 1a for schematic of region labels.

**FIG. S4.**
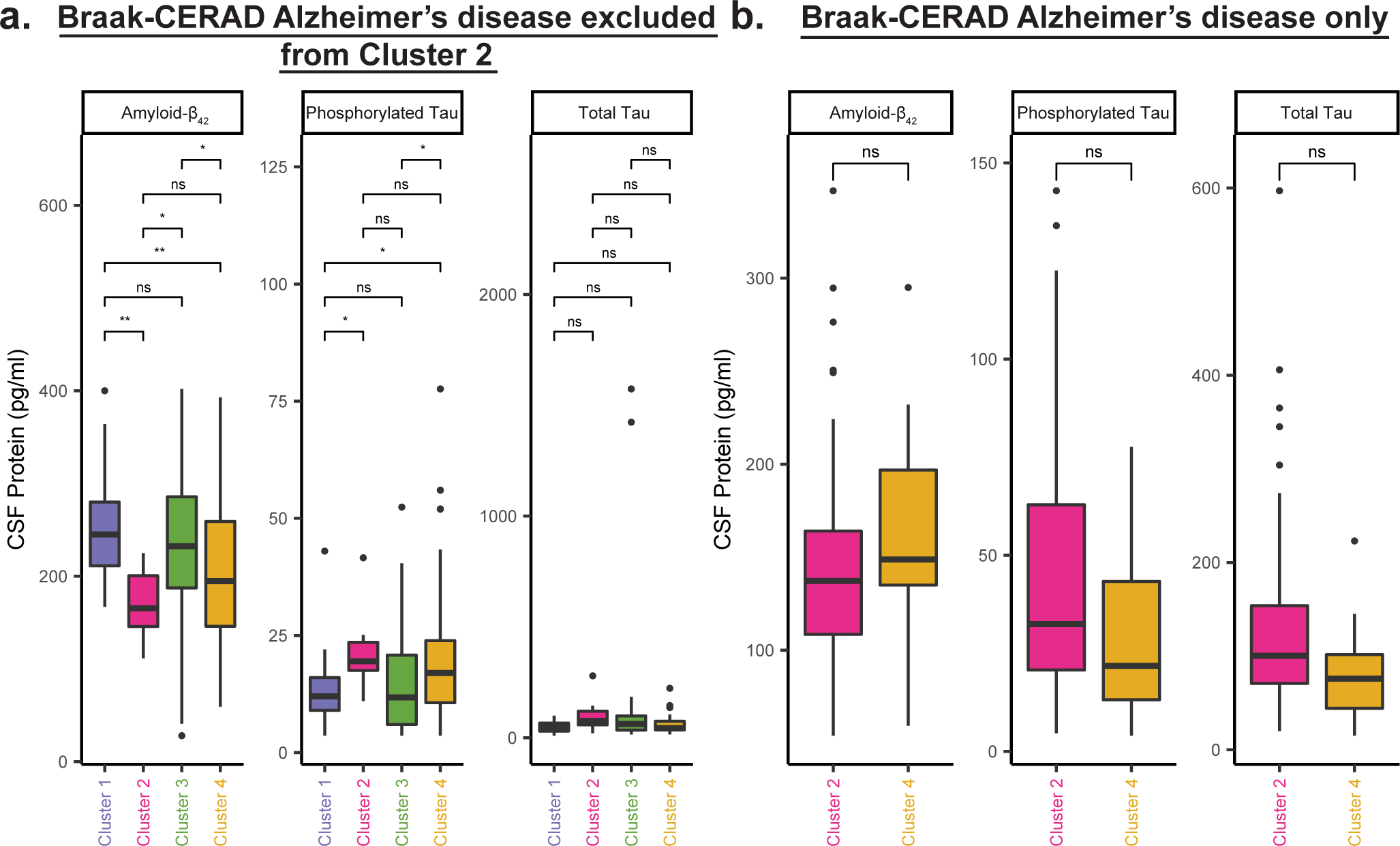
Differences in CSF protein levels between clusters extend beyond existing disease labels. *(a-b)* Pairwise intercluster comparisons of median CSF protein levels for amyloid-*β*_1−42_, phosphorylated tau, and total tau using Wilcoxon rank-sum test, FDR-corrected for multiple comparisons (*q <* 0.05) over all 3 proteins. In panel *a*, analysis is performed on an *n* = 126 subsample of patients with Braak-CERAD Alzheimer’s disease excluded from Cluster 2. In panel *b*, analysis is performed on an *n* = 159 subsample of patients, all of whom meet criteria for Braak-CERAD Alzheimer’s disease. Clusters 1 and 3 were excluded because they contained 2 or fewer patients with Braak-CERAD Alzheimer’s disease. ns, *p*_FDR_ *>* 0.05. *, *p*_FDR_ *<* 0.05. **, *p*_FDR_ *<* 0.01. ***, *p*_FDR_ *<* 0.001. ****, *p*_FDR_ *<* 10^*−*6^. CSF, cerebrospinal fluid.

### Variations on disease label prediction task

In the main text, we present prediction of cluster membership using an unpenalized multiple logistic regression classifier trained only on CSF protein levels. However, we attempted 3 variations on this approach in the experimental process. Conversion to a one-versus-all prediction, class-balancing, splitting of training and testing samples, and evaluation of out-of-sample performance was performed here exactly as in the main text (see Methods for details).

First, we trained an unpenalized multiple logistic regression model to predict membership to existing disease labels and disease clusters using both CSF protein and genotype at the *APOE* and *MAPT* loci. Compared to the out-of-sample accuracy and AUC achieved in Fig. 5 using only CSF protein, the performance of these models was virtually identical. Mean AUCs in predicting LBD, FTLD, and Alzheimer’s disease were 0.71 (Fig. S7a 95% CI, 0.70-0.72; Fig. 5a 95% CI, 0.69-0.71), 0.74 (Fig. S7a 95% CI, 0.73-0.74; Fig. 5a 95% CI, 0.74-0.76), and 0.94 (Fig. S7a 95% CI, 0.94-0.94; Fig. 5a 95% CI, 0.93-0.94), respectively. Mean AUCs in predicting Cluster 1, Cluster 2, Cluster 3, and Cluster 4 were 0.67 (Fig. S7b 95% CI, 0.67-0.68; Fig. 5c 95% CI, 0.67-0.69), 0.83 (Fig. S7b 95% CI, 0.82-0.83; Fig. 5c 95% CI, 0.83-0.84), 0.91 (Fig. S7b 95% CI, 0.91-0.91; Fig. 5c 95% CI, 0.91-0.91), and 0.84 (Fig. S7b 95% CI, 0.84-0.84; Fig. 5c 95% CI, 0.86-0.86) respectively. These findings suggest that genotype at the *APOE* and *MAPT* loci do not explain meaningful unique variance in disease class membership, under the traditional system of disease labels or our system of disease clusters.

We also used a random forest algorithm to predict disease class labels from both CSF protein and genotype at the *APOE* and *MAPT* loci, in case there were nonlinear relationships or interactions between genes, CSF, and class labels. This approach again produced similar performance to that of a multiple logistic regression model trained on CSF only. Prediction of Cluster 1 (Fig. S8c 95% CI, 0.88-0.89; Fig. 5c 95% CI, 0.86-0.86) and AD (Fig. S8a 95% CI, 0.91-0.91; Fig. 5a 95% CI, 0.93-0.94) may have been slightly more accurate, but prediction of Cluster 3 (Fig. S8c 95% CI, 0.80-0.81; Fig. 5c 95% CI, 0.83-0.84) was slightly less accurate. These findings suggest that accounting for non-linearities and interactions does not improve the ability to map phenotype and genotype at 2 loci to disease labels, and again supports the idea that genotype at the *APOE* and *MAPT* loci do not provide useful information beyond what can be obtained through CSF protein analysis.

**FIG. S5.**
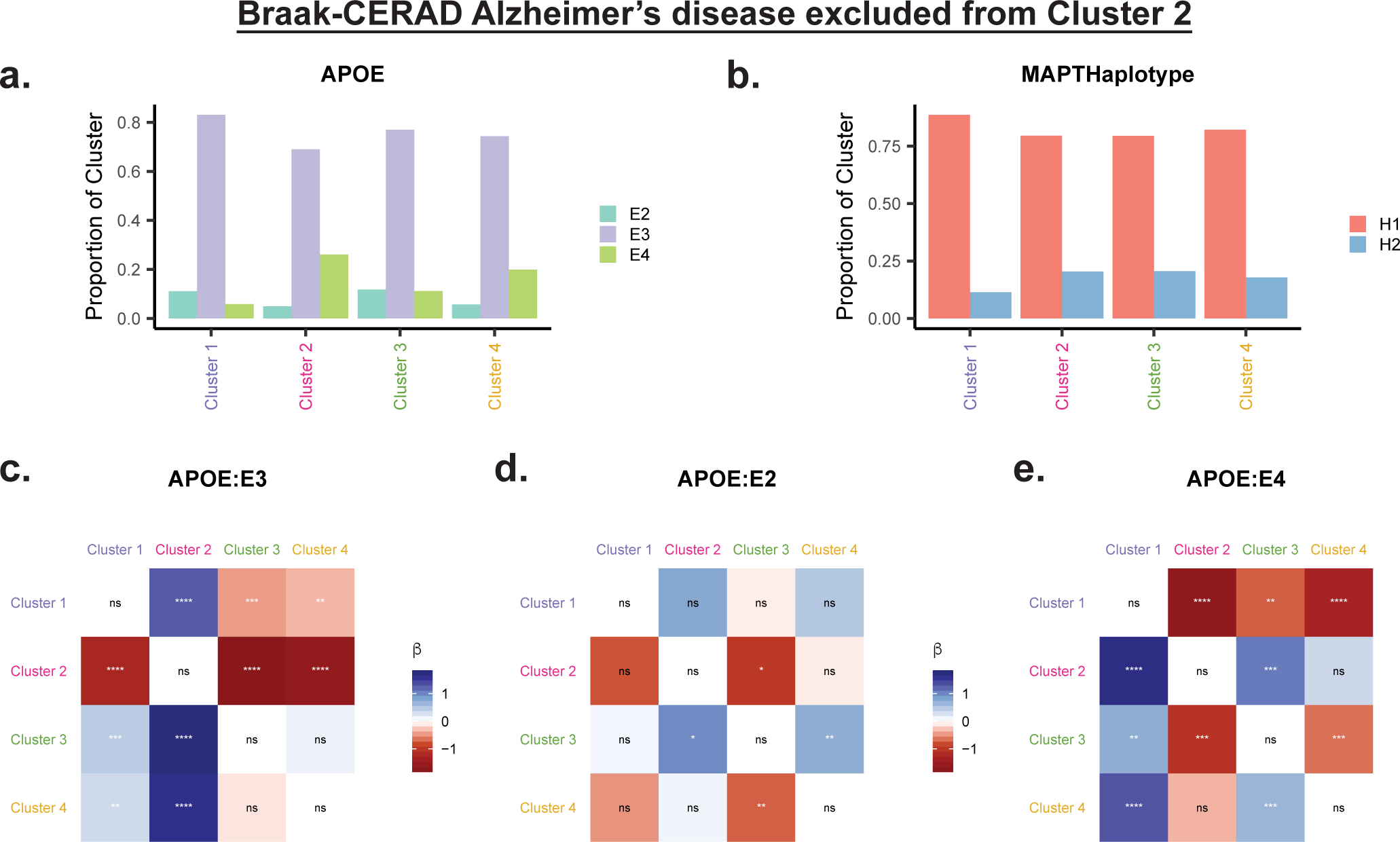
Genotypic differences between clusters at the *APOE* locus persist when excluding Alzheimer’s disease. Using an *n* = 887 sample excluding patients with Braak-CERAD Alzheimer’s disease from Cluster 2, we assessed the prevalence of *APOE* alleles across clusters. *(a-b)* Within each cluster, we calculated the proportion of each allele for *APOE (panel a)* and *MAPT (panel b). (c-e)* Matrix of logistic regression *β*-weights, whose *ij*^th^ element reflects the increase in log odds ratio for membership to cluster *i* relative to cluster *j* given the presence of *APOE* -*ϵ*2 *(panel c), APOE* -*ϵ*3 *(panel d), APOE* -*ϵ*4 *(panel e)*. ns, *p*_FDR_ *>* 0.05. *, *p*_FDR_ *<* 0.05. **, *p*_FDR_ *<* 0.01. ***, *p*_FDR_ *<* 0.001. ****, *p*_FDR_ *<* 10^*−*6^.

However, the fact that the genotypes we studied do not improve prediction in a model that already contains CSF protein data does not mean that prediction based on genotype is not potentially useful. In fact, prediction based on genotype may be more useful in that it is less invasive and could provide an estimate of risk for a particularly disease class before the onset of subclinical neurodegeneration. To assess this possibility, we trained a multiple logistic regression classifier to predict disease labels based on *APOE* and *MAPT* genotype. The classifier was able to predict PSP, Multiple System Atrophy, LBD, FTLD, corticobasal degeneration, amyotrophic lateral sclerosis, and Alzheimer’s disease with above-chance out-of-sample accuracy. Additionally, Cluster 1, Cluster 2, and Cluster 3 could be predicted out-of-sample with above-chance accuracy. Prediction of Alzheimer’s disease (Fig. S10a; 95% CI 0.71-0.71) and Cluster 2 (Fig. S10c; 95% CI 0.70-0.71) were very similar, consistent with the fact that these histopathologically defined groups overlap genetically (Fig. S5c-e and Fig. 4d-f).

Finally, we wished to assess the extent to which the strong representation of Alzheimer’s disease in Cluster 2 was driving the high classification accuracy of Cluster 2. We trained multiple logistic regression classifiers to predict membership to disease clusters from CSF protein levels after excluding Braak-CERAD Alzheimer’s disease from Cluster 2. We found that out-of-sample accuracy was diminished for all clusters, likely due in part to reduced sample size and the elimination of Alzheimer’s patients from Cluster 2, which represents a population of phenotypically distinct individuals. Nevertheless, we were able to identify Cluster 2 with AUC of 0.74. Overall, these results suggest that individuals in Cluster 2 without Alzheimer’s disease are somewhat phenotypically distinct from other clusters in terms of CSF protein levels, although not as much as individuals with Alzheimer’s disease.

**FIG. S6.**
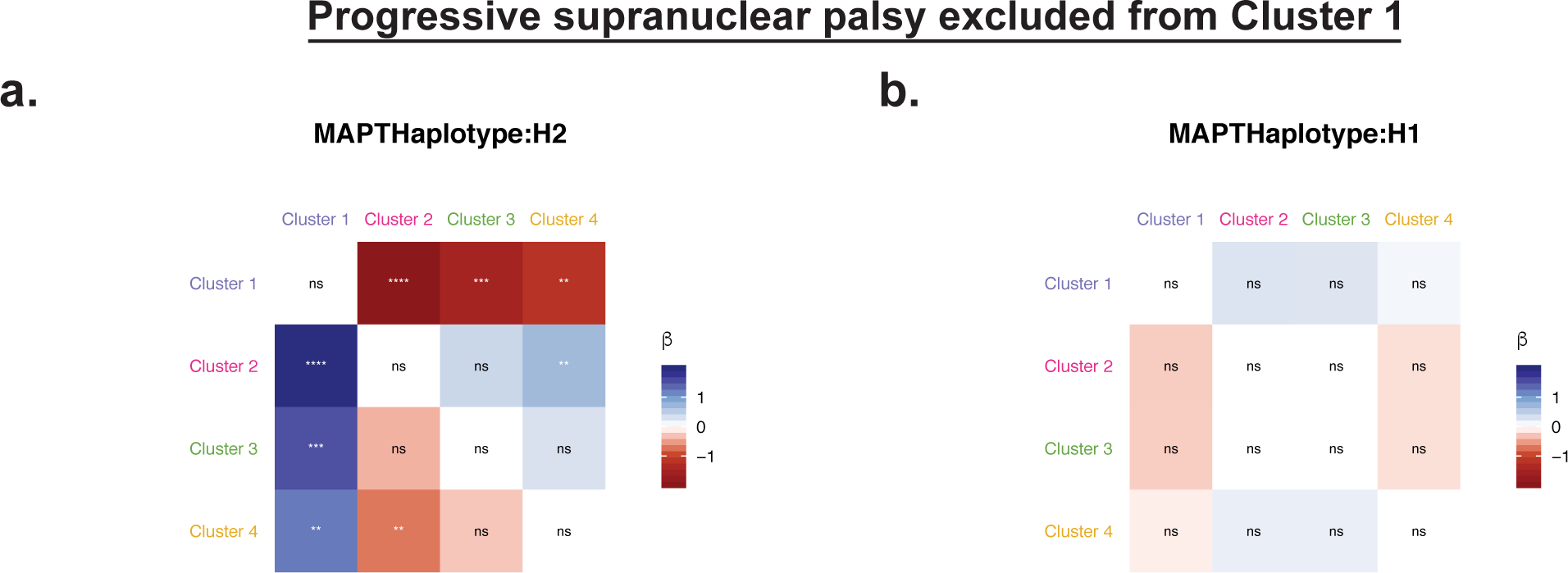
Genotypic differences between clusters at the *MAPT* locus persist when excluding progressive supranuclear palsy. In panels *a-b*, we analyze an *n* = 1239 sample excluding all patients with progressive supranuclear palsy. *(a-b)* Matrices of logistic regression *β*-weights, whose *ij*^th^ element reflects the increase in log odds ratio for membership to cluster *i* relative to cluster *j* given the presence of *MAPT* -H2 *(panel a)* or *MAPT* -H1 *(panel b)*. ns, *p*_FDR_ *>* 0.05. *, *p*_FDR_ *<* 0.05. **, *p*_FDR_ *<* 0.01. ***, *p*_FDR_ *<* 0.001. ****, *p*_FDR_ *<* 10^*−*6^.

**FIG. S7.**
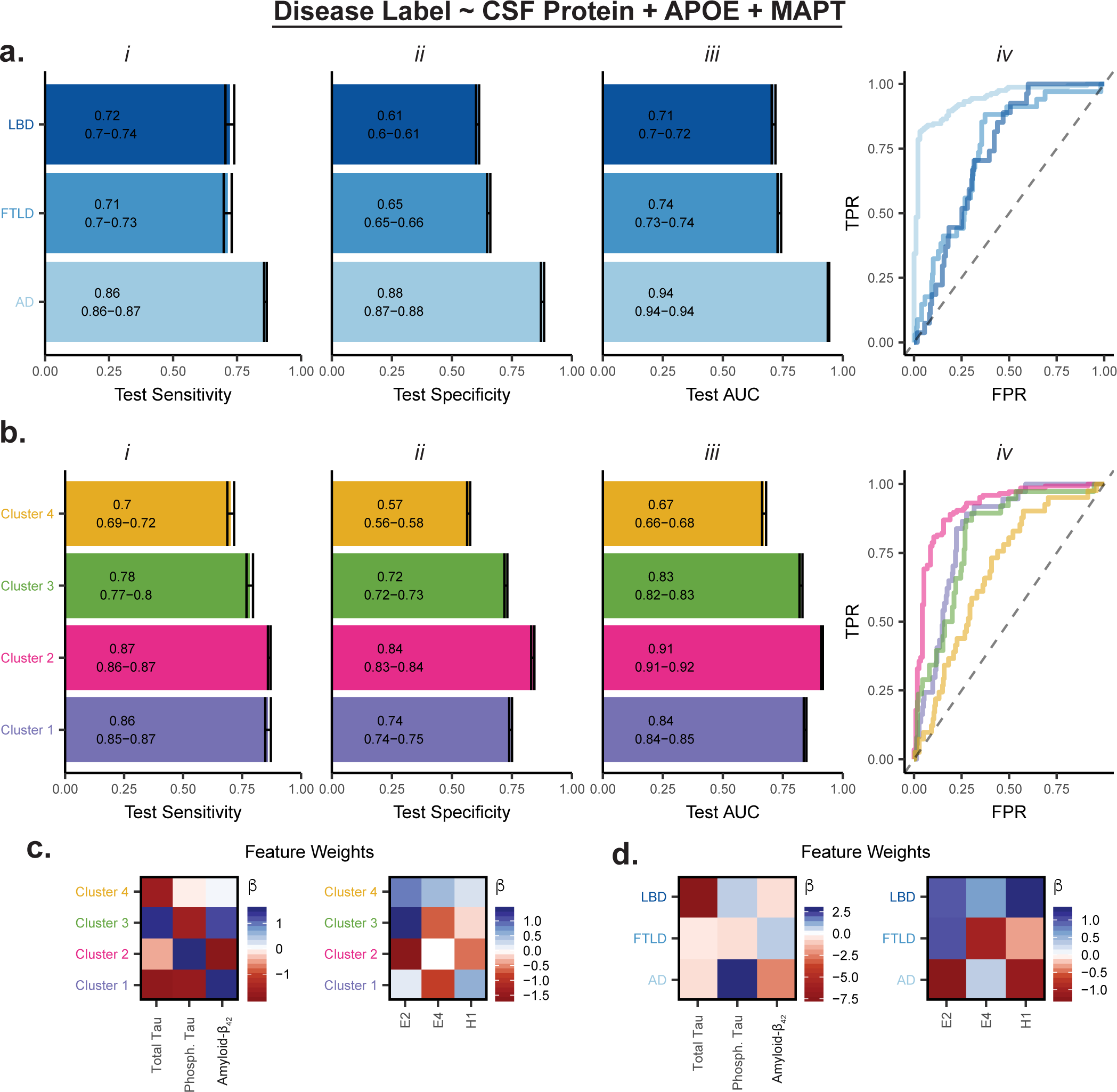
Genotype adds minimally to identification of disease labels when using CSF protein data. *(a-b)* Characteristics of prediction of existing diagnoses *(panel a)* or disease clusters *(panel b)* in held-out testing data using multiple logistic regression to predict disease labels from CSF protein levels and genotype. Sub-panels *i*, and *ii* show the test-set sensitivity and specificity, respectively, using a threshold value of 0.5. Sub-panel *iii* shows the area under the curve (AUC) on the test-set, reflecting performance over a range of threshold values. Bar length represents mean performance, and error bars indicate 95% confidence intervals over 100 repetitions of *k*-fold cross-validation at *k* = 10. Sub-panel *iv* shows representative receiver-operator characteristic curves for test-set predictions of existing diagnoses *(panel a)* or disease clusters *(panel b). (c-d)* Mean standardized multiple logistic regression *β* across 100 repetitions of *k*-fold cross-validation at *k* = 10 in the prediction task for existing disease labels *(panel e)* or in the prediction task for disease clusters *(panel f)*. The *β* weights can be interpreted as the increase in log-odds ratio for a one standard deviation increase in the value of the predictor in the case of continuous predictors, or for a one unit increase in allele count in the case of interval predictors. TPR, true positive rate. FPR, false positive rate. Total Tau, total CSF tau protein. Phosph. Tau, total CSF phosphorylated tau. Amyloid-*β*_1−42_, total CSF amyloid-*β*_1−42_. ϵ2 and ϵ4, *c*2 and *c*4 alleles at the *APOE* locus. H1, *MAPT* haplotype.

**FIG. S8.**
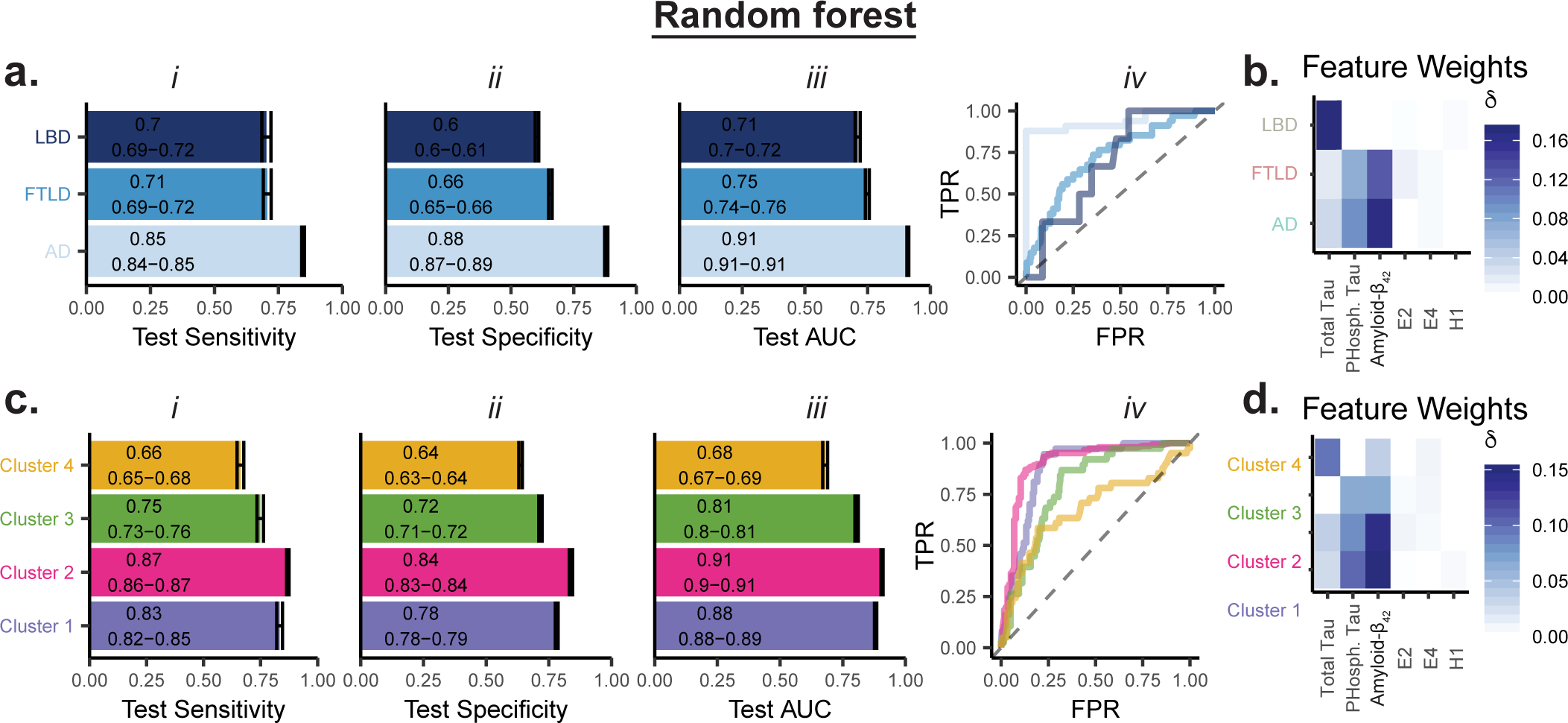
Random-forest algorithm exhibits similar performance to multiple logistic regression in identifying disease labels. *(a, c)* Characteristics of prediction of existing diagnoses *(panel a)* or disease clusters *(panel c)* in held-out testing data using a random forest classifier to predict disease labels from CSF protein levels. Sub-panels *i* and *ii* show the test-set sensitivity and specificity, respectively, using a threshold value of 0.5. Sub-panel *iii* shows the area under the curve (AUC) on the test-set, reflecting performance over a range of threshold values. Bar length represents mean performance, and error bars indicate 95% confidence intervals over 100 repetitions of *k*-fold cross-validation at *k* = 10. Sub-panel *iv* shows representative receiver-operator characteristic curves for test-set predictions of existing diagnoses *(panel a)* or disease clusters *(panel c). (b, d)* Representative decrease in accuracy *δ* when removing each predictor individually in the prediction task for existing disease labels *(panel e)* or in the prediction task for disease clusters *(panel f)*. A larger decrease in accuracy indicates that a feature was more important for a particular prediction task. TPR, true positive rate. FPR, false positive rate. Total Tau, total CSF tau protein. Phosph. Tau, total CSF phosphorylated tau. Amyloid-*β*_1−42_, total CSF amyloid-*β*_1−42_. E2 and E4, ϵ2 and ϵ4 alleles at the *APOE* locus. H1, *MAPT* haplotype.

**FIG. S9.**
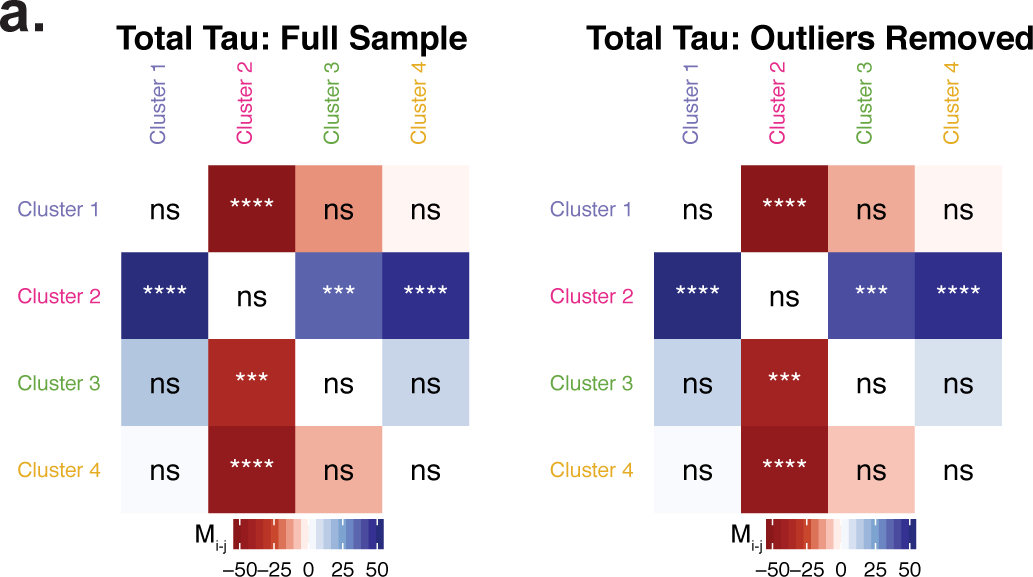
Differences in CSF total tau levels are robust to outliers. *(a)* Effect sizes and statistical significance of pairwise intercluster comparisons of median CSF total tau levels using Wilcoxon rank-sum test, FDR-corrected for multiple comparisons (*q <* 0.05) over all 3 proteins. In the left panel, the analysis is performed on the full sample as in Fig. 3b, and in the right panel, two outliers in Cluster 3 with total tau *>* 1000 pg/ml were removed. Results were virtually identical with or without these outliers included in the analysis. ns, *p*_FDR_ *>* 0.05. *, *p*_FDR_ *<* 0.05. **, *p*_FDR_ *<* 0.01. ***, *p*_FDR_ *<* 0.001. ****, *p*_FDR_ *<* 10^*−*6^. CSF, cerebrospinal fluid.

**FIG. S10.**
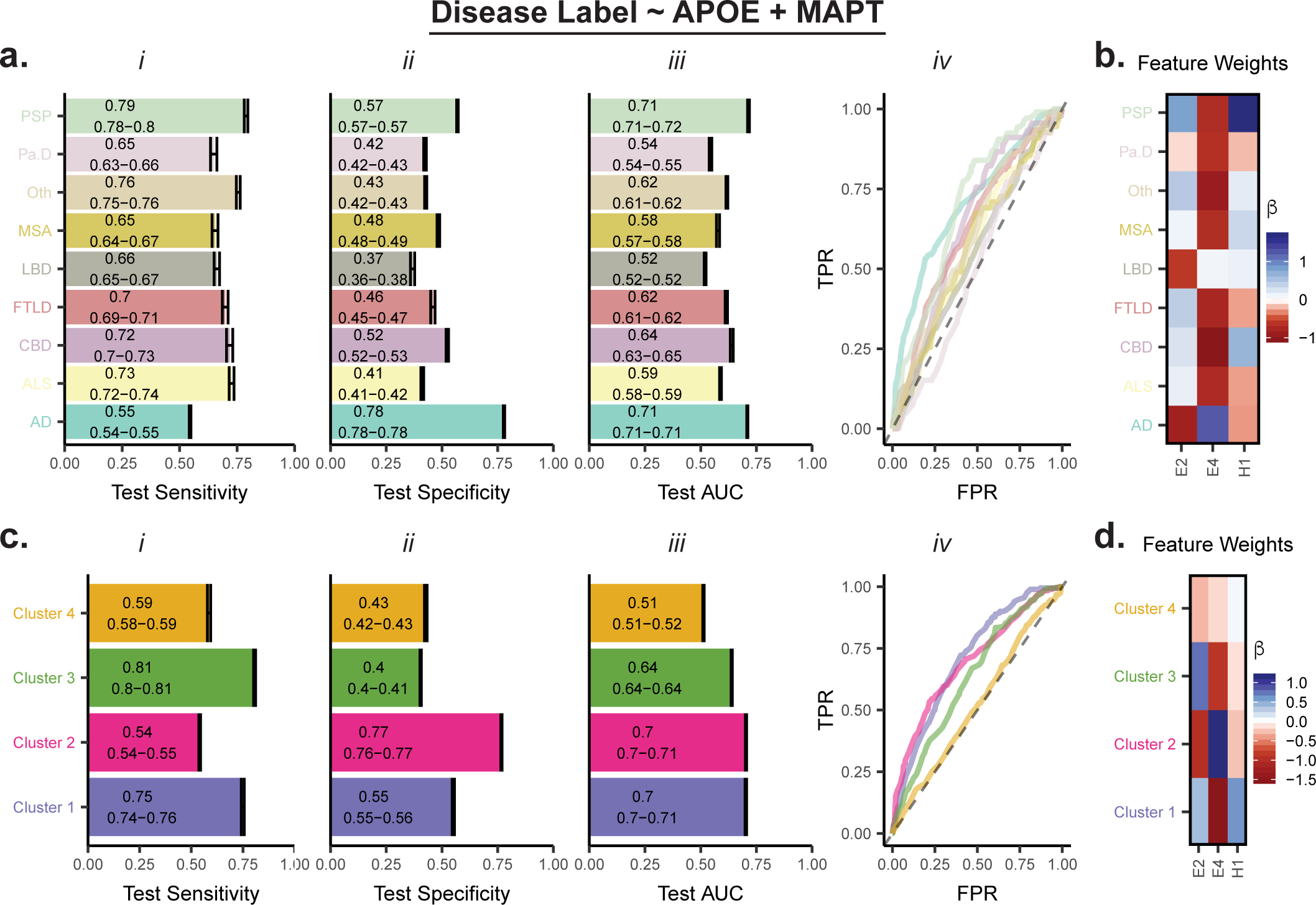
Disease labels can be predicted with above-chance accuracy from *APOE* and *MAPT* genotype. *(a, c)* Characteristics of prediction of existing diagnoses *(panel a)* or disease clusters *(panel c)* in held-out testing data using multiple logistic regression to predict disease labels from genotype. Sub-panels *i* and *ii* show the test-set sensitivity and specificity, respectively, using a threshold value of 0.5. Sub-panel *iii* shows the area under the curve (AUC) on the test-set, reflecting performance over a range of threshold values. Bar length represents mean performance, and error bars indicate 95% confidence intervals over 100 repetitions of *k*-fold cross-validation at *k* = 10. Sub-panel *iv* shows representative receiver-operator characteristic curves for test-set predictions of existing diagnoses *(panel a)* or disease clusters *(panel c). (b, d)* Mean standardized multiple logistic regression *β* across 100 repetitions of *k*-fold cross-validation at *k* = 10 in prediction task for existing disease labels *(panel e)* or disease clusters *(panel f)*. The *β* weights can be interpreted as the increase in log-odds ratio for a one unit increase in allele count. TPR, true positive rate. FPR, false positive rate. E2 and E4, *ϵ*2 and *ϵ*4 alleles at the *APOE* locus. H1, *MAPT* haplotype.

**FIG. S11.**
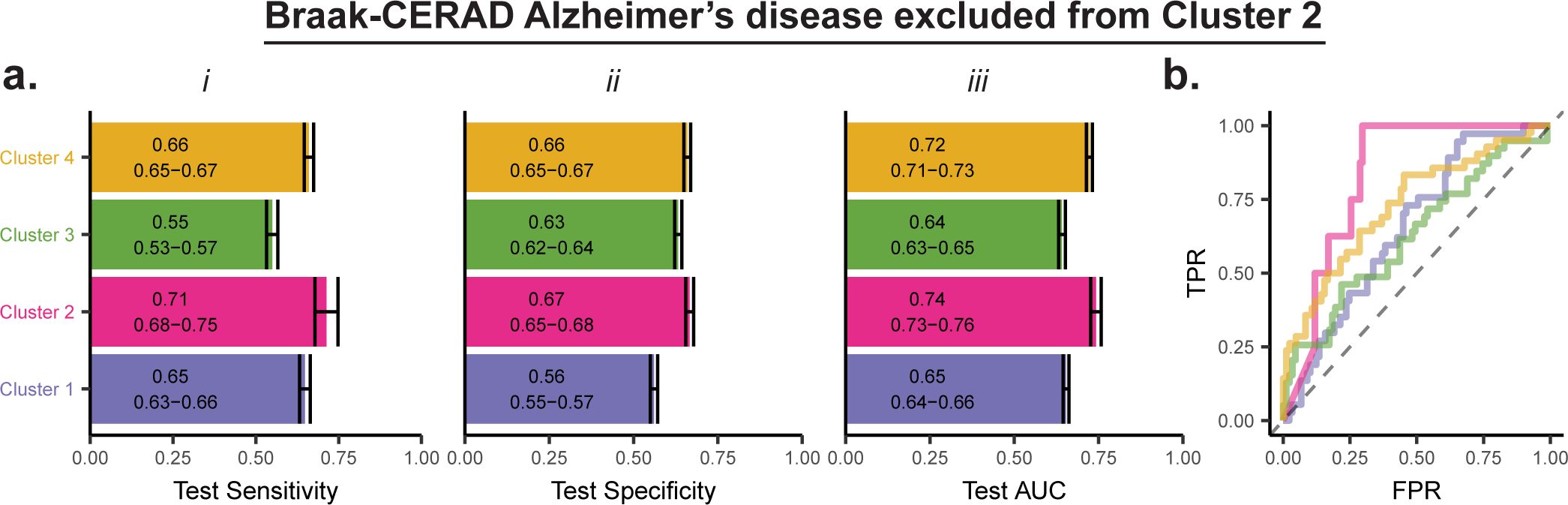
Non-Alzheimer’s disease patients in Cluster 2 can be identified with accuracy above chance. *(a)* Characteristics of test set prediction of disease clusters from CSF protein in an *n* = 126 sample with Braak-CERAD Alzheimer’s disease excluded from Cluster 2 only, using multiple logistic regression. Sub-panels *i* and *ii* show the test-set sensitivity and specificity, respectively, using a threshold value of 0.5. Sub-panel *iv* shows the area under the curve (AUC) on the test-set, reflecting performance over a range of threshold values. Bar length represents mean performance, and error bars indicate 95% confidence intervals over 100 repetitions of *k*-fold cross-validation at *k* = 10. *(b)* Representative receiver-operator characteristic curves for test-set prediction of disease clusters. TPR, true positive rate. FPR, false positive rate. Total Tau, total CSF tau protein. Phosph. Tau, total CSF phosphorylated tau. Amyloid-*β*_1−42_, total CSF amyloid-*β*_1−42_.

